# EasyAb: A High-Throughput Workflow for Antibody-Based PTM Peptide Enrichment Method Coupled to Mass Spectrometry

**DOI:** 10.1101/2024.10.29.620939

**Authors:** Ashok Kumar Jayavelu, Jeffrey J Liu, Sonja C Schriever, Paul T Pfluger, Christoph Schwarzer, Natalie Krahmer, Matthias Mann

## Abstract

Reversible post-translational modification (PTMs) is a fundamental mechanism of cellular signal transduction. *In vivo* and *ex vivo* studies to profile PTMs have greatly advanced our understanding of the complexities of cellular signaling. However, apart from the most commonly studied PTMs, large-scale analysis is still very challenging, limiting our understanding of various cellular processes. PTM-bearing peptides often enriched by antibodies, followed by unbiased mass spectrometry (MS)-based readout of hundreds or thousands of sites. To extend the reach of this powerful technology to in vivo and ex vivo studies with small protein starting amounts, we here developed EasyAb, a streamlined and high throughput MS workflow for antibody-based PTM profiling. Using epidermal growth factor receptor (EGFR) signaling and acute myeloid leukemia (AML) cell systems, we demonstrate that EasyAb increases sensitivity and enables multiple large-scale systems-level studies. Furthermore, EasyAb resolves in vivo brain G protein-coupled receptor-mediated tyrosine kinase activation and reveals the long elusive hypothalamic neuron-specific leptin signaling architecture.

**Graphical Abstract:** 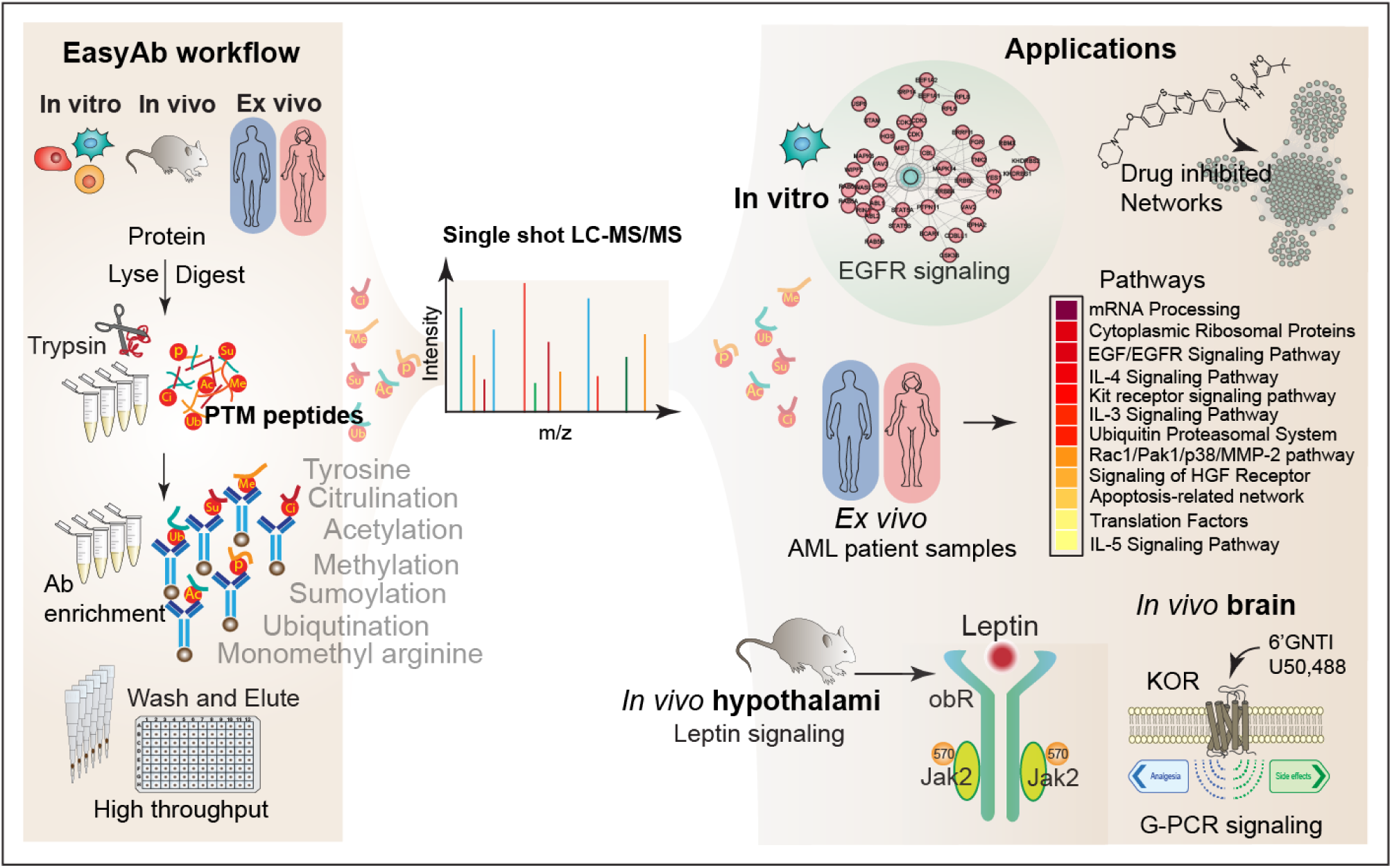

**HIGHLIGHTS:** EasyAb, a sensitive, rapid and high-throughput method to quantify post translationally modified peptides in diverse cell systems.

EasyAb reveals differential tyrosine phosphorylation of mRNA splicing proteins in AML patient samples. Activation of KOR by “G-protein-biased” non-aversive agonist 6’GNTI elicits Src kinase activity in mice. Elucidation of *in vivo* hypothalamic Leptin induced LEPRb-Jak2 phosphotyrosine signaling architecture.

## INTRODUCTION

Protein post translational modification events such as acetylation, methylation, ubiquitination, sumoylation, tyrosine phosphorylation are key spatial and temporal controls of cellular processes including signaling transduction (Choudhary and Mann, 2010; Deribe et al., 2010). PTM of proteins lead to rapid changes in their activity, interaction, and localization. This is especially important for the control of cellular functions that needs to respond to abrupt fluctuations of nutrient levels on a seconds-scale to daily basis. (Humphrey et al., 2015). Despite their crucial function, there are relatively few comprehensive PTM profiling studies available from tissues (Bennett et al., 2007; Hoffman et al., 2015; Labots et al., 2017; Lundby et al., 2012; Sathe et al., 2018; Yi et al., 2018), which is mainly due to limited amounts of sample and lack of sensitive sample preparation workflows. Improving on these limitations would have major implication in our basic understanding of disease such as cancer, diabetes, obesity etc. that are driven by complex PTM network changes (Fan et al., 2015; Humphrey *et al*., 2015). As a prominent example of interest to our groups, the landscape of leptin regulated phosphotyrosine changes at the hypothalamic arcuate nuclei is central in obesity but is still not defined due to technological challenges. Identification of leptin regulated novel targets at the hypothalamus could shed light on key players in energy metabolism in the brain, potentially contributing to new therapeutically relevant approaches. More generally, insights into the dysregulated complex PTM networks of disease such as diabetes, obesity, cancer and drug abuse from animals and human samples are invaluable to understand the underlying cellular processes involved.

Advances in mass spectrometry (MS)-based proteomics can now quantify PTMs at increasing depth and speed, enabling unprecedented systems-level views (Bekker-Jensen et al., 2020; Hogrebe et al., 2018). We previously described EasyPhos, a high-throughput TiO2 based phosphopeptide enrichment method, which allowed elucidation of systems-wide and temporally resolved *in vivo* action of insulin in liver tissues (Hoffman et al., 2015; Humphrey et al., 2015; Humphrey et al., 2018). However, the liver is a relatively large organ and advances such as µPhos workflow can now enable to assess phosphoproteome of small organs or tissues invitro and *in vivo* (Oliinyk et al., 2024) . Furthermore, phospho-serine and phospho-threonine can be efficiently captured by chemical matrices such as TiO2 (Humphrey *et al*., 2018; Larsen et al., 2005), but most other proteomic PTM workflows typically require complex and cumbersome antibody (Ab) enrichment of the PTM-containing peptides prior to MS analysis. This often demands high protein input (generally 10-20 mg) especially for label free quantification approaches, requires multiple buffer exchanges, takes several days (Abe et al., 2017a; Rikova et al., 2007; van der Mijn et al., 2015; Wu et al., 2015; Xu et al., 2010), and often results in significant amount of peptide loss and inconsistent data quantification (**Figure S1A-C**). This constitutes a barrier to the application of Ab-based PTM proteomics, especially to animal or clinical studies for signaling pathway elucidation, targeted drug development, novel biomarkers identification, and drug response monitoring.

The workflows used so far applied conventional cell lysis and digestion protocols with sodium dodecyl sulphate (SDS), urea, or guanidinium hydrochloride (Gdmcl) buffers which tend to be laborious and preclude direct coupling of digestion to enrichment (Lundby et al., 2019; Scholz et al., 2015). Specifically, the incompatibility of lysis (irrespective of label free or chemical labelling approaches), digestion and enrichment buffers currently necessitate multiple buffer exchanges that introduce sample loss, variation in quantitation and extra handling steps. To overcome these challenges, we here develop EasyAb, an approach that permits rapid (one-day), cost-effective, and in-depth quantification of PTM bearing peptides in a high-throughput manner. Our one-step digestion-to-enrichment workflow circumvents multiple buffer exchange, time consuming and expensive strategies involving peptide fractionation or multiplexing the samples by chemical labelling to boost peptide identification (Gajadhar et al., 2015; Lamoliatte et al., 2014; Rose et al., 2016; Udeshi et al., 2013; Zhang et al., 2010). Our strategy collapses many steps of earlier workflows, limiting sample loss and enabling high-throughput *in vivo* and *ex vivo* applications in order to gain insights into biological processes that were previously inaccessible (**Figure 1B**).

**Figure 1.**
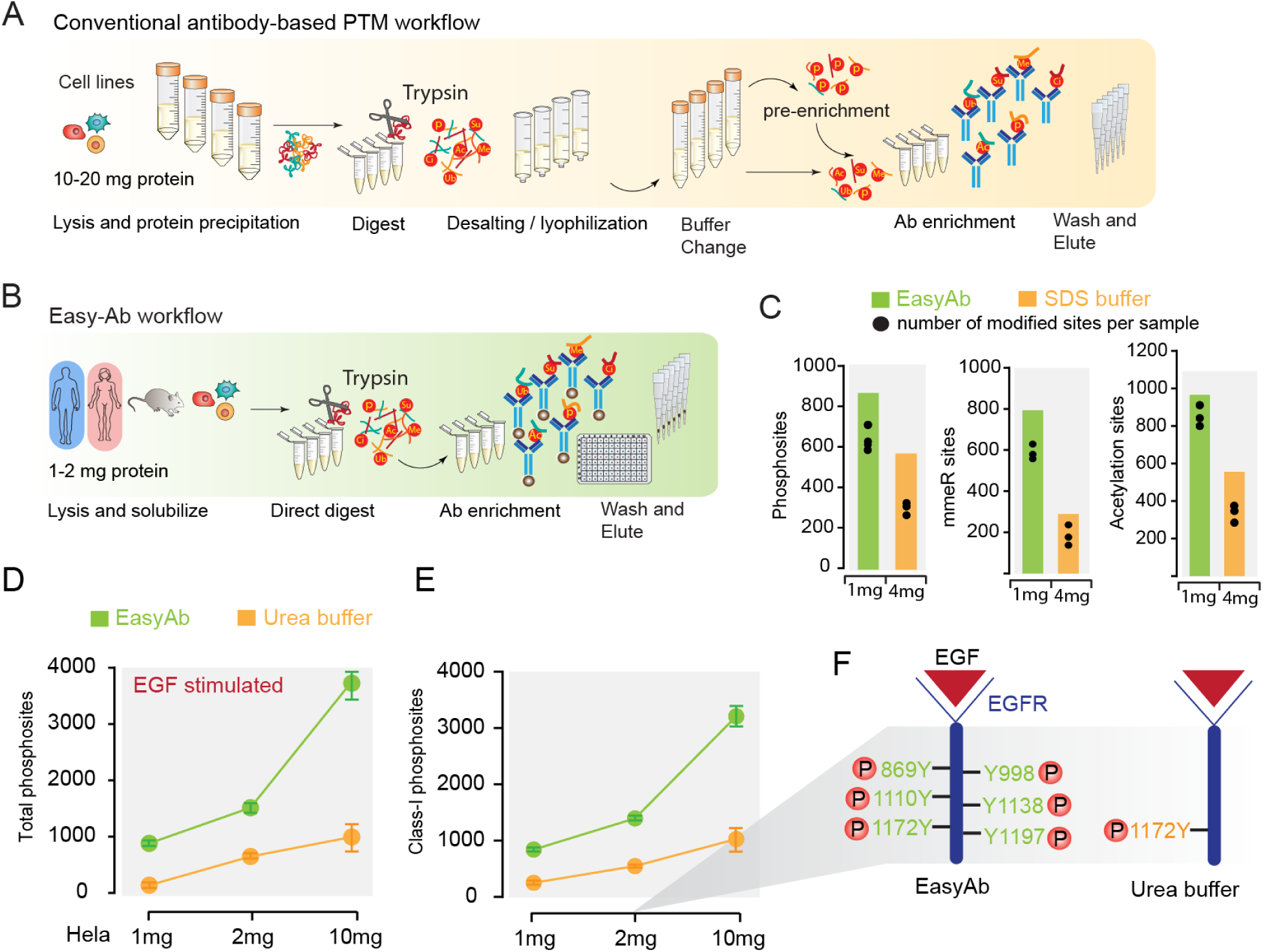
Benchmarking EasyAb against conventional workflows. (**A**) Overview of conventional antibody-based PTM enrichment workflows. (**B**) Overview of EasyAb workflow highlighting low sample input, reduced steps and high throughput. (**C**) Comparison of EasyAb with the conventional SDS buffer workflow. The bar plot shows total number of identified sites in each experiment in a combined analysis and aligned dot plot shows number of identified sites per sample. Gain in combined analysis is the performance of match between runs (MBR) in transferring identification of phosphopeptides and quantification between the replicates in MaxQuant. The data shows EasyAb with 1 mg compared to conventional SDS buffer workflow with 4mg of starting material of HeLa lysate. For phosphotyrosine peptides (n=3-4, using pY1000 antibody), acetylated lysine (n=3, using AcK-100 antibody) and monomethyl arginine peptides (n=3, using memR antibody). (**D-F**) Comparison of EasyAb with Urea buffer workflow in HeLa cells stimulated with EGF. (**D**) The dot plot shows number of identified phosphosites at various amount of EGF stimulated Hela using various amounts of starting material (n=2, represent biological replicates). For comparison of 2mg data see Supplementary Table S1. (**E**) Represents the class-I phosphosites (high confidence phosphosites with localization probability score >0.75) identified (n=2, represent biological replicates). (**F**) Identified EGFR phosphotyrosine sites from 2mg starting material in EasyAb vs Urea buffer.

In proof-of-principle experiments we show EasyAb applications with antibodies targeting acetylation, monomethyl arginine, and tyrosine phosphorylated peptides. We benchmark EasyAb sensitivity using the well-characterized EFG stimulated EGFR receptor signaling in mammalian cells and demonstrate the superiority of EasyAb in comparison to existing workflows. We display the utility of the higher throughput of EasyAb in diverse biological areas such as time resolved analysis of EGFR activated targets and in the elucidation of phosphotyrosine signatures in patient derived AML cells lines for identification of rewired kinases, drug targets, and pathway dependencies. Furthermore, to validate the *ex vivo* application of EasyAb we explore the landscape of FLT3 receptor signaling in AML patient samples. Using limiting amount of mouse brain samples, we resolve *in vivo* brain G protein-coupled receptor (KOR)-mediated tyrosine kinase activation upon treatment with aversion and non-aversion agonists. Finally, and most importantly, we elucidate the hypothalamic neuron-specific leptin signaling architecture, by producing the first *in vivo* phosphotyrosine leptin signaling map.

## RESULTS

### Development and benchmarking of the EasyAb workflow

Conventional Urea buffer and SDS buffer-based lysis demands protein precipitation and desalting prior to enrichment of PTM peptides, which contributes to 30-40% loss of sample amounts (Supplement figure 1A-C). Sodium deoxycholate (SDC) based buffers are compatible with both lysis and protease digestion conditions (Lin et al., 2008; Yu et al., 2003). It enhances the activity of trypsin, increases the number of protein identification and facilitates the recovery of hydrophobic peptides compared to other digestion buffers such as SDS (Lin et al., 2013; Masuda et al., 2008). Thus they eliminate the need for protein precipitation or desalting chromatography, one of the major steps of protein loss (Humphrey *et al*., 2018; Kulak et al., 2014; Lin *et al*., 2008; Liu et al., 2018; Zhou et al., 2006). Conventional immunoprecipitation buffers such as RIPA contains SDC; however, the buffer composition is not compatible with efficient protease digestion (due presence of NP40 and Triton X) and requires buffer exchanges and several washing steps to remove detergents prior to MS (Scheerlinck et al., 2015). Further significant loss of protein amount occurs, we questioned whether with adjustment of SDC concentration and the pH of the lysate post protease enzymatic digestion, the same buffer would be amenable to Ab-enrichment as well, thus eliminating the need for desalting and buffer exchanges (see **Figure S1D**-**G,** for development and optimization. Of note, the PTM antibodies binds to modified sites by recognizing motifs on protein and peptide). Furthermore, we wanted to establish a simplified protocol that can be easily adapted to a high throughput 96 well format (**Methods**). Here we optimized a one-step digestion-to-enrichment workflow and named it EasyAb. Using this simplified protocol dramatically reduces the required starting material, the time required for sample processing to less than one day, and is highly cost effective. With antibodies conjugated to magnetic beads EasyAb is adaptable to a 96-well plate format for parallel processing and reduction of experimental error.

As proof-of-principle experiments, we benchmarked our developed EasyAb workflow against the conventional ones using 4%SDS or 6M Urea buffer for sample amounts and sensitivity. We preformed enrichment with well validated PTM antibodies targeting acetylation (AcK), monomethyl arginine (mem-R), and tyrosine phosphorylated (pY1000) in unperturbed Hela cell lysates prepared by either EasyAb or SDS workflow, and analyzed the modified peptides (at a false discovery rate (FDR) less than 1% in this study, Method section) by single run LC-MS/MS. We consistently found increased identification of post-translationally modified peptides in samples prepared by EasyAb when compared to samples prepared with SDS buffer (**Figure 1C**). Notably this was detected using four-times less sample input amount (1mg vs 4mg). Thus, EasyAb should be generally applied to antibody based PTM workflows.

We next benchmarked EasyAb to Urea buffer workflow by comparing the tyrosine peptide enrichment depth using various amounts of EGF stimulated Hela cell lysates. Here again we detected many fold increase in phosphotyrosine peptides using EasyAb compared to Urea workflow (**Figure 1D-E**). Moreover, similar depth and sensitivity was achieved with fivefold less starting material compared to the conventional Urea method, which requires 10 mg for similar results. We highlighted the EGF receptor phosphorylation as an example of the sensitivity of EasyAb (**Figure 1F and Table S1**). This gain in the depth of coverage does not attribute to the impact of new series of Q-Exactive platforms since the MS samples prepared by conventional protocol were measured with the same setting and were still out performed by the sample prepared using the EasyAb workflow. Thus, the EasyAb protocol produces uncompromised depth of modified peptide coverage with several-fold reduced starting material and higher reproducibility between replicates (**Figure S1H-I**; R^2^= 0.87-0.98) compared to published protocols, suggesting that EasyAb should greatly improve performance of antibody-based enrichment (for details on the further optimization see **Supplementary Information**) prior to MS measurement.

### EasyAb enables in-depth tyrosine phosphoproteome analysis

Tyrosine phosphorylation comprises only 0.5-1% of total phosphorylation events in cells but plays a disproportionally important regulatory role in signal transduction (Hunter, 2014). Antibody-based enrichment of tyrosine phosphopeptides from large amounts of cell or tissue samples has to almost universally employed to examine the tyrosine phosphoproteome (Lundby *et al*., 2019; van der Mijn *et al*., 2015). We applied our Easy-Ab workflow using anti-phosphotyrosine antibodies to diverse biological systems to examine its versatility *in vitro, in vivo* and ex *vivo* experiments and to gain insights into biological systems that were previously inaccessible.

First, we performed label free quantification in the AML cell lines Molm13 and MV4-11 harboring receptor FLT3 kinase internal tandem duplication mutation (FLT3-ITD). From just 2 mg starting material, we identified 4,416 phosphosites (**Figure S2A-B**; for details see **Supplementary Information**) with high reproducibility (FDR of less than 1%, Pearson correlation R^2^ between 0.78-0.92) (**Figure S2F**). In comparison to EasyAb, the protocol from others (Abe et al., 2017b; Labots *et al*., 2017; Tong et al., 2017) and our previous studies (Sharma et al., 2014), took 4-5 days instead of one day; and in the latter case-at four or more times the material - only achieved half the phosphopeptide coverage. Other protocols in the literature required even larger amounts of materials and employed strategies like multiplexing sample materials through chemical labeling, pre or post-fractionation to achieve less than half of these numbers (Abe *et al*., 2017b; Batth et al., 2018; Bian et al., 2016; Boersema et al., 2010; Fang et al., 2020; Tong *et al*., 2017; van Alphen et al., 2020; van der Mijn *et al*., 2015; Zhang *et al*., 2010). Our deep coverage illuminates some previously less noticed aspects of this well studied system. For example, characterization of membrane protein PTMs has been more challenging due to issues in solubilization and enrichment (Orsburn et al., 2011). Using EasyAb, over 40% of the quantified proteins were membrane or transmembrane proteins, with 30% of the sites previously unreported (**Figure S2H**). We identified seven and quantified five phosphotyrosine sites in the FLT3 receptor (**Figure S2I**), including a novel pY401 site. Together, the FLT3 mutated cell presents phosphorylation of 31 tyrosine kinases and 14 phosphatases (**Figure S2J**), of which many of them (Btk, Syk, Abl, Src, and Ptpn11) were known target of FLT3-ITD signaling.

To illustrate the utility of EasyAb in quickly uncovering biological and drug related insights, we studied the Quizartinib-mediated response of mutant FLT3-ITD kinase receptor, a proven AML drug target (Daver et al., 2019). Quizartinib (**Figure S2K**), a potent and highly specific mutant FLT3 kinase inhibitor, is promising in treating adults relapsed/refractory AML with the FLT3-ITD mutation (Cortes et al., 2019) and we therefore examined Quizartinib induced global phosphotyrosine changes in the corresponding cell lines. In line with previous observations Quizartinib treatment significantly (t-test analysis, permutation based FDR <0.05) decreased phosphorylation levels of FLT3-ITD kinase (Zarrinkar et al., 2009) and its downstream targets Stat5 and Mapk (Hou et al., 2017). The inhibitor also dramatically downregulated phosphorylation on well-characterized targets of FLT3-ITD such as Syk, Btk, Gab2, Grb2, Ptpn11 (**Figure S2L**) thus attenuating FLT3-ITD kinase signaling. Pathway analysis of significantly regulated phosphoproteins reveals enrichment of “metabolism”, “apoptosis”, “response to stress”, and “cell cycle” highlighting the major cellular pathways rewired by FLT3-ITD (**Figure S2M**). At the site level we observed dynamically decreased phosphorylation of Gsk3A (Y279) and Lair1 (Y281), revealing critical inhibitory mechanisms of apoptosis hijacked by FLT3-ITD kinase (**Figure S2L**). The deep phosphoproteome also revealed novel links to TNK2 and STAT6 signaling (additional details in **Supplementary Information** and **Figure S3**) and uncovered new Quizartinib-responsive phosphosites that underlie pathway rewiring.

### EasyAb facilitates time-resolved EGFR signaling

We next assessed whether EasyAb is suitable for time-resolved analysis with reduced sample amounts. To demonstrate this on an extensively studied system (Blagoev et al., 2004; Fang *et al*., 2020; van der Mijn *et al*., 2015), we stimulated Hela cells from a single petri dish (<2 mg protein) with EGF (50ng/ml) for 0, 15 or 60 minutes (**Figure S4A**). We observed phosphorylation on known targets of EGFR signaling including Y727, Y998, Y1110, and Y1172 on EGFR and phosphorylation of Met, Ptpn11, Gsk3b and Stat5 (**Figure S4B**). Furthermore, we identified > 1000 tyrosine sites at the FDR of <1% and after filtering for quantified phospho S/T/Ysites, multiple sample t-test analysis (permutation-based FDR <0.05) revealed dynamic phosphorylation changes on 84 phosphosites (**Figure S4C**). This included EGFR phosphotyrosine sites and several novel substrates regulated by EGF stimulation such as Vav2 (Y437), Lars (Y939), Fbxl7 (Y354), and Dock11 (Y1830) (**Figure S4D-E**). Thus, the EasyAb workflow enables in-depth and time resolved quantitative comparison of biological samples in a label free system.

### Application of EasyAb in *ex vivo* patient samples and patient-derived cell lines

To test if EasyAb is applicable to investigation of patient materials and compatible with large-scale analysis, we performed phosphotyrosine profiling of AML patient samples (n=14, **Table S2**) and 10 patient-derived cell lines harboring distinct driver mutations (n=3, per line as biological triplicates, **Figure S5A**) from 2 mg starting material. In total, this resulted in the identification of 10,013 phospho-S/T/Ysites (including singly, doubly, and greater phosphopeptides species), ∼80% of which were phosphotyrosine sites ( all class-I sites, localization probability >0.70). This is, the largest number of phosphotyrosine sites identified in a single cell type of myeloid origin under native condition so far (**Figure 2A**). Remarkably, about 1770 tyrosine sites were novel and about 18% lack known sequence motifs (**Figure S5 C,D**). Furthermore, 36% of total phosphorylation events were on membrane proteins (**Figure S5E**).

**Figure 2.**
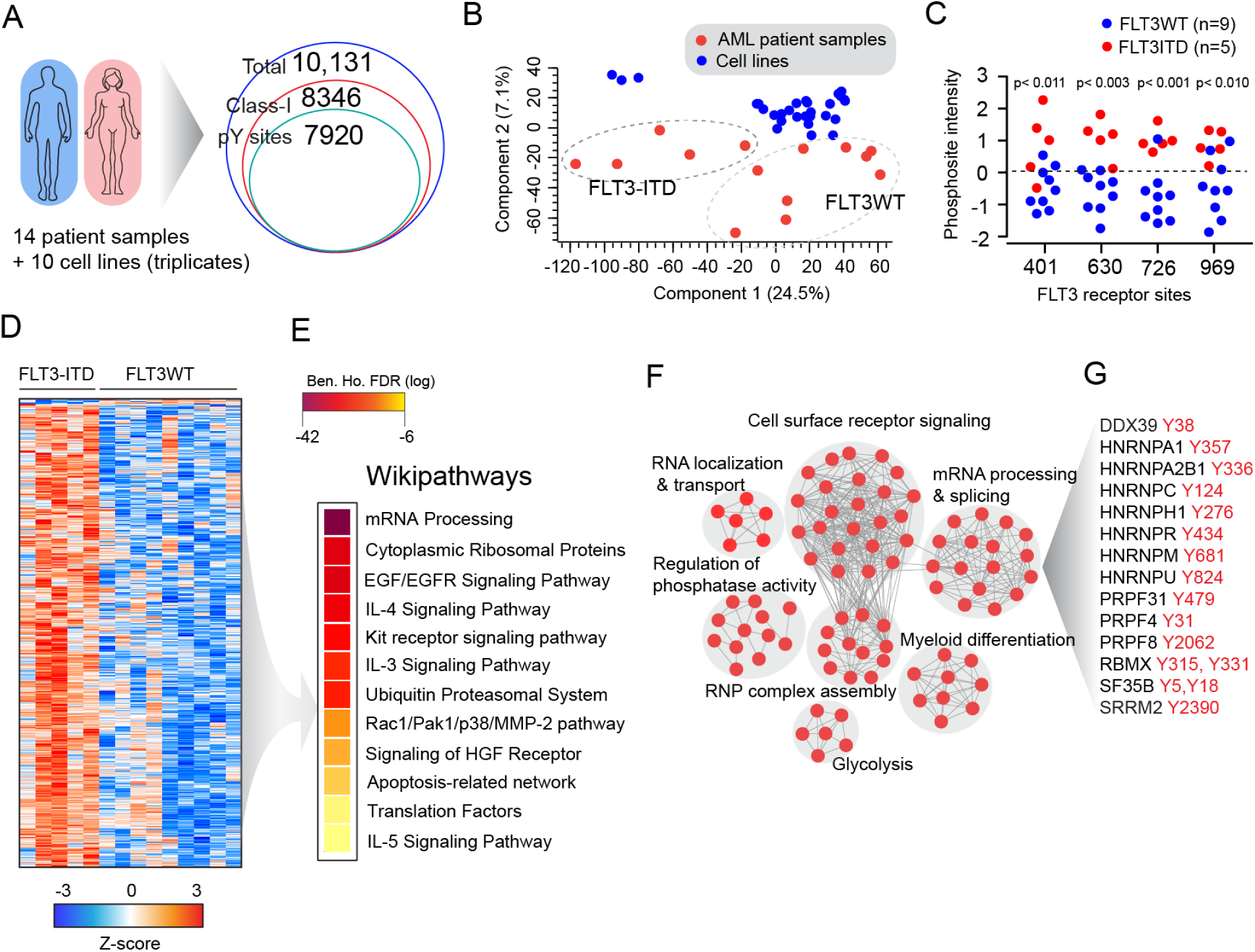
*Ex vivo* AML patient phosphotyrosine signaling using EasyAb. (**A**) Number of total, class-I, and phosphotyrosine sites identified in AML patients sample (n=14) and patient derived leukemic cell lines (n=10) from 2mg sample material. See also Figure S5. (**B**) Principle component analysis (PCA) shows clustering of the AML cell lines and patient samples based on the phosphotyrosine phosphoproteome. (**C**) Scatter dot blot shows the regulated FLT3 receptor phosphorylation sites in AML patient samples grouped based on the genotype, FLT3WT (n=9) and FLT3-ITD (n=5). (**D**) Heatmap of the significantly regulated phosphosites in AML patient samples based on their FLT3 status. (**E**) Heatmap of enriched wikipathways in AML with FLT3-ITD. (**F**) Cytoscape network map displays the GO terms enriched from significantly tyrosine phosphorylated proteins in FLT3-ITD AML. (**G**) Core mRNA splicing factors and its tyrosine phosphosites significantly regulated in AML with FLT3-ITD.

In AML patient samples alone, we identified a staggering 6,339 phosphotyrosine sites and quantified nearly 2300 sites (**Figure 5B and Figure S5F**). Principal component analysis (PCA) revealed a clear separation of cell lines and patient samples based on their phosphotyrosine profile (**Figure 2B**). This supports the notion that cell lines which are frequently used as in vitro models, do not entirely reflect the biology of primary patient samples, emphasizing the need for investigation of actionable drug targets directly on clinical samples. To this end, our AML patient sample cohort consisted of two prognostic genotypes FLT3WT (n=9) and FLT3ITD (n=5). As expected, greater number of phosphosites were quantified in mutant FLT3 receptor tyrosine kinase samples (constitutively active state) compared to wild type receptor (endogenous inactive, unstimulated state; **Figure S5F**). Two sample t-test analysis (FLT3WT vs FLT3ITD, **Table S3**) revealed regulation of >500 sites (permutation-based FDR<5%), including FLT3 receptor tyrosine phosphorylation (**Figure 2C**). Several known signaling molecules among these includes STAT5, Syk, Btk, Gab2, Grb2, Ptpn11 as identified in the above FLT3-ITD cells lines. In addition, we identified differential tyrosine phosphorylation of 8 autophagy (e.g. ATG2B), 24 apoptosis, 30 cell cycle, 54 cell surface proteins (e.g. CD97, CD99L2), 13 DNA damage & repair proteins, and 36 kinases (e.g. PTK2B, TYK2) in FLT3-ITD patients. Perhaps most interestingly, one of the key enriched pathways was ‘mRNA processing & splicing’ in this dataset (**Figure 2E**). In many cancers including hematological malignancies splicing factors are commonly mutated leading to aberrant splicing that contributes to disease development and progression (Lee and Abdel-Wahab, 2016). Here we discover differential phosphotyrosine regulation of mRNA splicing proteins (**Figure 2F,G**) driven by an oncokinase. The functional significance of this novel regulatory mechanism in driving the cancer would be an interesting window into novel signaling mechanisms to explore.

The AML cell lines fell into three main clusters, indicating common footprints of phosphotyrosine signatures between them (**Figure S6A-C**). Fisher’s exact test on kinase substrate motifs revealed strongest enrichment for Src kinase substrate motifs in cluster III followed by cluster II and cluster I (**Figure S6D**). Enriched Src kinase motifs were independent of Src tyrosine kinase expression levels suggesting that the activity of Src kinases is crucial. To validate this hypothesis, we treated the different clusters of cell lines with the Src kinase inhibitor Bosutinib, which confirmed that the cluster-III cell line MM6 was most sensitive to the Src kinase inhibitor (**Figure S6E**). Conversely, the profiled cell lines harbor specific mutations, which could potentially drive novel tyrosine signaling. Mutation-specific changes included several known and novel key signaling molecules such as PDGFR (Y894), BTK (Y551), INNP5D (Y1022), and TAOK3 (Y429) tyrosine phosphorylated specific to cell lines (**Figure S7,** for additional details and further characterization of the cell lines see **Supplementary Information**).

### In vivo brain GPCR phosphotyrosine signaling

To demonstrate *in vivo* applications, we next explored phosphotyrosine signaling in the brain. G protein-coupled receptor (GPCRs) signaling, such as kappa opioid receptor (KOR) mediate neurotransmitter metabotropic signaling is essential in neuronal plasticity and functions. Activation of KOR by “G-protein-biased” agonists alleviates pain without eliciting unwanted aversion or hallucination (White et al., 2015). We recently dissected this signaling system is a large-scale study resulting in the temporal and spatial distribution of tens of thousands of phosphosites. However, only about a hundred of these were pY sites (Liu *et al*., 2018). Here, we applied EasyAb to tissues obtained from cortex, hippocampus, striatum, and medulla oblongata of a single animal (number of experimental animals, n=3 per group), which was sacrificed 5 minutes after administration of the saline control, U50,488H the aversive agonist, or 6’GNTI - the non-aversive agonist. Cortex had the most tyrosine phosphorylation sites and other regions progressively less in the order of cortex > medulla oblongata > hippocampus > striatum (**Figure 3A**). PCA of each brain region revealed their own unique tyrosine phosphoproteomic signatures, much like their counterparts in the general phosphoproteome and proteome (Liu *et al*., 2018; Sharma et al., 2015). Although brain region-specific tyrosine kinase expression level showed similar trend (**Figure 3B,C**), the expression levels of these kinases no longer correlated with the linear motifs of the most abundant tyrosine phosphopeptides in each region, further demonstrating that tyrosine kinases are more closely regulated than serine and threonine kinases in tissues. In addition, we observed region-specific perturbation of the tyrosine phosphoproteome by the aversive agonist U50, 488H that paralleled those of the general phosphoproteome (**Figure 3D**). In the cortex, the aversive agent U50, 488H and non-aversive one 6’GNTI each induced unique perturbations (ANOVA-significant) of the tyrosine phosphoproteome from the saline-treatment mice (**Figure 3E**). Linear kinase motifs of Jak2 and Src were both enriched in the ANOVA-significant sites (**Figure 3F**) with phosphopeptides containing Src motifs contributing to more than 40% of the total. This agrees with our previous observation that Src is dephosphorylated by U50, 488H but not 6’GNTI treatment (Liu *et al*., 2018). Fischer’s exact test of Src kinase substrates reveals categorical enrichment of both ‘ionotropic glutamate receptor activity’ and ‘coated vesicle’, indicating that the aversive KOR-mediated inactivation of Src influences glutamate signaling and neurotransmitter release through pY based mechanisms. To further validate this finding, we inhibited Src in vivo using PP2, a selective Src family kinases inhibitor (n=3 mice per group, 5mg/kg, 30 mins before saline or 6’GNTI). This confirmed that the non-aversive agonist 6’GNTI elicits Src kinase activity which is prevented upon PP2 treatment (**Figure 3G**).

**Figure 3.**
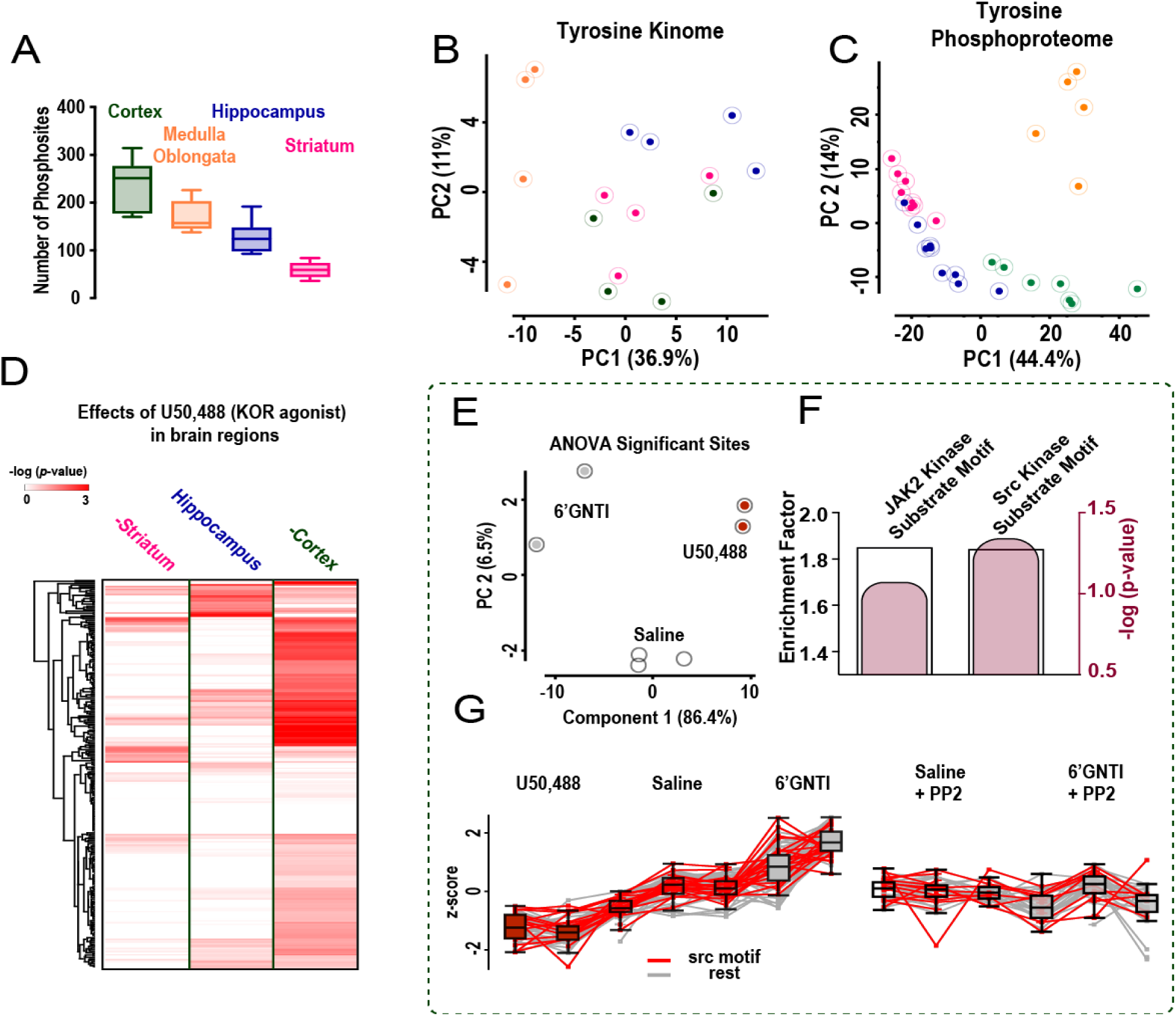
GPCR mediated phosphotyrosine signaling on various brain regions of mice. (**A**) Boxplot showing the number of phosphotyrosine sites quantified in each brain region. (**B**) PCA of tyrosine kinases found in the relevant brain regions (color-coded according to Figure.3A). (**C**) PCA of tyrosine phosphorylation sites in the relevant brain regions (color-coded according to Figure.3A). (**D**) Effect of a reference kappa opioid receptor (KOR) agonist (U50,488) on the relevant brain regions visualized by heat map of p-values derived from pair-wise *t*-test between U50,488 and saline treated mice; number of changes is greater in cortex so that the ligand comparison was performed using samples from cortex. (**E**) PCA of ANOVA significant sites in mouse brain cortex; U50,488 (n=2) KOR agonist that induce aversion and 6’GNTI (n=2) is the KOR agonist that does not induce aversion. (**F**) Fischer exact test analysis of ANOVA significant sites, the black bar indicates the enrichment factor, whereas the pink bar indicates –log (p-value). (**G**) The profile plot of ANOVA significant sites; the red traces indicate regulated phosphosites with Src kinase substrate motif after treatment with U50,488, 6’GNTI (left panel) and after Src kinase inhibitor PP2 treatment (right panel).

In addition to Src substrates, we also found changes of phosphorylation states of the kinases Gsk3a/b (Y279/Y216), Dyrk2 (Y380) and Dyrk3 (Y368) (**Figure S8A**). Gsk3b is particularly interesting because it is involved in regulation of mTOR signaling a key finding in our previous study (Liu *et al*., 2018). The aversive agonist-mediated de-phosphorylation of the Gsk3a/b autophosphorylation site discovered here suggests that Gsk3b potentiates mTOR-mediated aversive signaling (Kitagishi et al., 2012), highlighting how EasyAb can illuminate signaling pathways in pharmacologically important systems.

### In vivo mouse hypothalamic leptin signaling

Obesity and diabetes are interlinked metabolic disease such that being obese vastly increases the chances of developing diabetes. Obesity involves impaired leptin hormone signaling, which is central in effectively regulating food intake and body weight. Leptin acts on the hypothalamic pro-opiomelanocortin (POMC) neurons of the central nervous system, however, despite its key roles the targets of activated leptin receptor signaling are still ill defined at the systems level. Leptin mediated signaling in the hypothalamic arcuate nucleus is even more challenging to elucidate, so that to our knowledge no systems-level signaling studies have been reported to date. Leptin activates the long form of the leptin receptor (LepRb), which is highly expressed in this major center governing food intake, body weight and energy metabolism. Activated LepRb signaling involves a cascade of known phosphotyrosine targets such as JAK2 tyrosine kinase and STAT3 (Banks et al., 2000). Here, we applied EasyAb to the hypothalamus of chow-fed C57Bl/6J mice that had received an intraperitoneal injection of vehicle or leptin (3 mg/kg body weight**, Figure S8C,D**). Subsequent phosphotyrosine and proteome profiling identified 639 pY sites on 173 proteins (overall identified phosphosites 1104 on 247 proteins); in total 271 phosphosites containing pS/T/Y were quantified and 65 pY sites were significantly regulated (permutation-based FDR <0.05. **Figure S8E,F and Table S4**).

We directly observed LEPRb-Jak2 activation by quantitative changes in known leptin target sites such as the autophosphorylation site Y570 of Jak2 kinase (Gong et al., 2007) and Y699 of STAT5b (**Figure 4**). We further quantified pY546 on Ptpn11, a protein tyrosine phosphatase (PTP) involved in the regulation of LepR-Jak2 signal transduction (Li and Friedman, 1999). In addition to these known PTPs, we discovered regulation of the receptor Ptprf at the Jak2 substrate-binding site on Y1381, suggesting that Ptprf may act as a negative regulator of LepR-Jak2 signaling. Furthermore, we quantified non-canonical members of leptin signaling such as Jnk1 (Y185) and Jnk3 (Y223), whose involvement in response to high glucose levels has been reported in endothelial cells (De Nigris et al., 2015).

**Figure 4.**
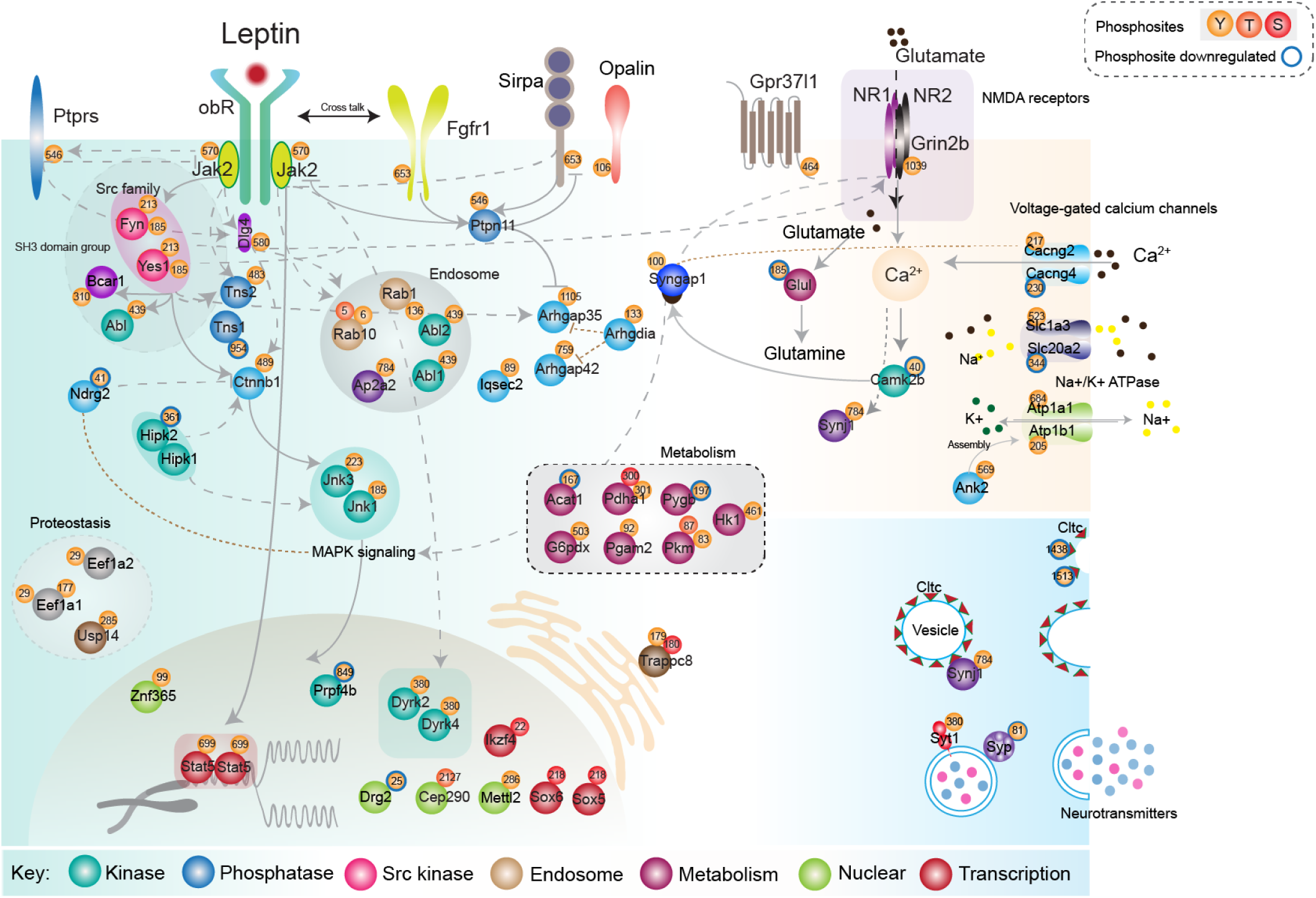
Leptin induced phosphotyrosine signaling network in the hypothalamus. Significantly regulated phosphosites in the mouse hypothalamus upon leptin treatment. The elucidated leptin map represents known canonical and non-canonical leptin signaling members and protein connections that are experimentally determined in the literature. Majority of the proteins in the network were organized based on their function. The circle on the protein denotes the phosphorylated residue (Tyrosine (Y), Threonine (T) and Serine (S) are represented in different colour as illustrated in the map). The blue ring around the circles of the phosphorylated residues = downregulation of phosphosite upon leptin action; no ring = upregulated upon leptin action. Established associations are connected in solid lines/arrows and curated associations are connected in dotted lines/arrows.

Our phosphotyrosine signaling network revealed several novel connections between leptin signaling and other major signaling networks. Activation of the non-receptor tyrosine protein kinase Fyn promotes Jak2 independent of STAT3 signaling upon leptin stimulation (Jiang et al., 2008). It is required in multiple cellular processes including brain development (Serpente et al., 1996) and axonal guidance (Morse et al., 1998), and is activated upon leptin stimulation at its autophosphorylation sites Y185 (Ferrando et al., 2012) and Y213 (Kaspar and Jaiswal, 2011). In line with this, we found regulation of Fyn at Y214 and Catenin beta-1 (Ctnnb1) at Y489, a component of canonical Wnt signaling and a known Fyn kinase target (Piedra et al., 2003). As the deletion of ß-catenin improves glucose tolerance and resistance to high fat diet (Elghazi et al., 2012), our data thus suggest an interplay between leptin-induced Fyn activation and beta-catenin Wnt signaling.

PSD95 was another interesting scaffold protein affected by leptin, phosphorylated on Y240 and Y580. We further observed leptin-simulated phosphorylation on PSD95 interacting partners Dlgap1 (Y75) and Dlgap3 (Y142 & Y723). PSD95 contains a PDZ domain along with SH3 and guanylate kinase domains that are often found in anchoring proteins, and act as a signal integrator. PSD95 is known to interact with NMDA glutamate receptors and GPCRs (Pedersen et al., 2017). Interestingly, PSD95 levels are known to be regulated by leptin and are implicated in neuronal signal transduction and brain development (Ramos-Lobo et al., 2019). We also observed leptin-induced phosphorylation of glutamate receptors Grin2b (Y1039) and Gpr37l1 (Y464) which are potential interacting partners of PSD95. Overall, these data suggest a trajectory of LEPRb-Jak2-driven phosphorylation events involving PSD95.

Another interesting connection is that of leptin with insulin and growth factor signaling. The membrane docking protein Sirpa is highly expressed in the brain and regulates growth factor as well as insulin tyrosine kinase receptor activity (Shen et al., 2009). Sirpa is a constitutive scaffolding protein for Jak2 (Galbaugh et al., 2010) and upon tyrosine phosphorylation binds to multiple proteins including protein tyrosine kinase Ptk2b, growth factor receptor-bound protein 2 (Grb2) and the PTPs Ptpn6 and Ptpn11(Shen et al., 2010). We found Sirpa to be significantly regulated at pY505 following leptin stimulation, likely due to Jak2 activation. Phosphorylated Sirpa recruits Ptpn11 in complex with Jak2 (Stofega et al., 2000) to induce phosphorylation of Ptpn11 at Y546, which could serve as a negative feedback mechanism through a gatekeeping role in the recruitment of Ptpn11 to the plasma membrane. Sirpa furthermore scaffolds with insulin like growth factor-1 to mediate Grb2-Pdk1 docking and Akt activation (Shen *et al*., 2010). Sirpa could thus serve as a potential link for an interaction of leptin and insulin signaling. Leptin-induced tensin 1 (Tns1) phosphorylation at Y954 and Tensin 2 (Tns2) phosphorylation at Y483 is another potential point of interplay between hypothalamic insulin and leptin action that emerges from our data. Functional tensin enzyme activity was shown to be regulated by Jak2, and is a prerequisite for the activation of Akt (Jung et al., 2011).

We also found increased phosphorylation of fibroblast growth factor 1 receptor (Fgfr1), a major regulator of metabolic homeostasis and insulin sensitization (Suh et al., 2014), at Y653. In rodents, the activation of hypothalamic FGF1R by a single intracerebroventricular injection of FGF1 was able to induce the sustained remission of diabetic hyperglycemia (Brown et al., 2019). The discovery of a leptin FGFR1 crosstalk mechanism within the hypothalamus is thus an interesting observation with relevance for both the CNS control of type 1 as well as type 2 diabetes (Jonker et al., 2012; Suh *et al*., 2014). We independently confirmed the above leptin regulated novel sites identified through MS by performing western and immunoprecipitation analysis on phosphotyrosine sites for which antibodies were available **(Figure S8G-I)**.

## DISCUSSION

Reversible PTM changes enable active switch of protein function to control essentially all cellular processes. The writers of PTMs such as kinases, methyltransferases, ubiquitin ligases, acetyltransferases, arginine methyltransferases are established therapeutic targets in cancer, metabolic diseases and several other disorders (Choudhary et al., 2014; Ferguson and Gray, 2018; Jarrold and Davies, 2019; Vucic et al., 2011). Given this importance, there has been relatively little focus on profiling these PTM modifications in human or animal samples, mainly due to lack of sensitive and rapid approaches. (Abe *et al*., 2017b; van der Mijn *et al*., 2015; Xu *et al*., 2010).

We here developed the EasyAb workflow and demonstrated that it is a rapid and sensitive tool with a number of key advantages. We successfully applied our protocol to the identification of peptides bearing diverse modifications such as acetylation, monomethyl arginine and phospho-tyrosine. EasyAb greatly decreases sample preparation time, reduces the amount of starting materials, is cost effective, and provides great depth of PTM coverage as benchmarked in EGFR signaling and AML cells lines. In higher throughput experiments, we show that EasyAb allows in-depth quantification of phosphotyrosine peptides from AML cell lines and *ex vivo* patient samples. In human AML patient samples, we identified the deepest tyrosine phosphoproteome to date. This revealed novel regulation of mRNA splicing factors in a subset of AML patient samples positive for FLT3 receptor mutation. We found activation of Src kinase and CDK in selected cell lines (described in the supplementary information, FigureS6 and FigureS7) and validated this finding using selective inhibitors, suggesting their therapeutic potential in AML. We further illustrate the sensitivity of our approach in our *in vivo* systems. This revealed novel signaling events involved in KOR-mediated aversive and non-aversive responses. Analyzing the mechanism of action of non-aversive agonist 6’GNTI on KOR unraveled the regulation of Src kinase and its substrates at the level of phosphotyrosine, which has eluded us before (Liu *et al*., 2018). The regulation of Src kinase by the GPCR agonist 6’GNTI was further confirmed by inhibition of Src kinase activity in vivo using PP2, a selective Src family kinase inhibitor.

Leptin activates the long form of the leptin receptor (LepRb), which is highly expressed in the hypothalamic arcuate nucleus, a major center governing food intake, body weight and energy metabolism. (Halaas et al., 1995) By applying the EasyAb workflow to hypothalami of mice treated with leptin, we confirmed known leptin signaling targets such as Jak2, STAT5 and identified numerous novel leptin regulated signaling components. Leptin induced effects extended to the non-canonical members Jnk, Src kinase Fyn, scoffold protein PSD95, and membrane docking protein Sirpa. In addition, novel leptin FGFR1 cross talk was discovered by EasyAb-MS. Notably, less than 10% of the regulated phosphoproteins had been annotated as part of the leptin pathway in the curated database Netpath (Kandasamy et al., 2010). Thus, our methods allowed us to generate the first large-scale map of leptin signaling and we achieved this in the *in vivo* situation from very limited input material. Our data reveal a plethora of previously unknown leptin mediated signaling events, and our atlas may serve as a useful resource to the metabolism research community (**Figure 4**). Future studies will be required to elucidate the functional roles of the new leptin targets in the leptin signaling network. It is intriguing that leptin signaling and KOR signaling activated some of the same kinase signaling pathways in different brain regions (Nicolas et al., 2013; Nicolas et al., 2012). These included the Jak2 mediated phosphorylation of Sirpa and Tns, the glutamate signaling pathways and components such as Fyn, NMDA receptor and PSD95. These results further indicate crosstalk or reuse of components in a general signaling network with tissue-specific modifications.

We envision that these advances in Ab-based PTM studies will open up further large-scale experiments on important signaling pathways in specific anatomical regions. While we have mainly focused on pY signaling, technologies like EasyAb can clearly provide unprecedented insights into any PTM that can efficiently be enriched by antibodies, providing crucial tools for the biological community to investigate complex PTM-regulated signaling networks.

## ACKNOWLEDGEMENTS

We thank all the members of Proteomics and Signal Transduction Department at the Max Planck Institute of Biochemistry, in particular Igor Paron, Christian Deiml, Bianca Splettstoesser, Dirk Wischnewski, Gaby Sowa, Alex Strasser for their technical assistance. We also thank Dr. Frank D Böhmer for the critical reading and comments on the FLT3-ITD signaling phosphoproteome. Medini Steger critically read and improved the manuscript. This work carried out for this manuscript was supported by the Max Planck Society for the Advancement of Science and by German Research Foundation (DFG/Gottfried Wilhelm Leibniz Prize). C.S. received grants P 30430 and P 30592 from the Austrian Science Fund. A.K.J is funded by Emmy-Noether DFG(JA3274/1-1). N.K. is funded by Emmy-Noether DFG (KR5166/1-1). S.C.C and P.T.P received supported from the German Center for Diabetes Research (DZD). P.T.P by the Helmholtz-Israel-Cooperation in Personalized Medicine and European Union within the scope of the European Research Council ERC-CoG Yoyo-LepReSens (no. 101002247).

## AUTHOR CONTRIBUTIONS

A.K.J and M.M conceived the project and wrote the manuscript with inputs from all authors. A.K.J developed the EasyAb workflow. A.K.J and J.J.L performed the MS experiments. A.K.J, J.J.L, S.C.S, P.T.P, C.S, N.K, and M.M designed, performed the animal experiments, interpreted the data, and performed computational and statistical analysis.

## STAR METHODS

### Cell lines

HeLa, cells were cultured in DMEM medium supplemented with 10%FBS and 1% Penicillin/Streptomycin (Pen/Strep). Molm13, MV4-11, HEL, EOL-1, Kasumi-1, KG1, OCI-AML3, SD-1, ML2, HL60 lines were maintained in RPMI 1640 supplemented with 10% heat inactivated FBS and 1% penicillin/streptomycin. MM6 cells were additionally supplemented with 2mM glutamate, 1mM sodium pyruvate and 10µg/ml insulin (Thermo Fischer Scientific). OCI-AML5 cell were grown in α-MEM supplemented with 20% heat inactivated FBS and 10ng/ml rhGMCSF (Immuno Tools). All the lines were maintained at 37°C under 5% CO2.

### Antibodies

For PTM-enrichment experiments for EasyAb, magnetic beads conjugated pY1000 antibody (Cat No 14017S), Acetylated-Lysine (Ac-K2-100) MultiMab^TM^ Rabbit mAb mix (Cat No 9814S) and Mono-Methylated Arginine (mme-R) MultiMab^TM^ Rabbit mAb mix (Cat No 8015S) were purchased from Cell Signaling Technology. Anti-phosphotyrosine 4G10 antibody was a gift from Dr.Frank D Böhmer ( generated in-house at CMB, University Hospital Jena). Secondary antibody for western blotting (goat-anti-rabbit HRP conjugate, Cat No NA934V) was purchased from Sigma.

### Sample preparation for the EasyAb Benchmark experiments

HeLa cells were grown to ∼70% confluence in DMEM medium supplemented with 10% FBS in 15 cm petri dishes. Cells were stimulated with 100 ng/ml of recombinant human epidermal growth factor (Cat No 11343406, ImmunoTools) in serum free DMEM condition for 20 minutes, washed with ice cold PBS, and lysed in SDC, Urea or SDS buffer. The SDC buffer contained 4% SDC, 100 mM Tris-HCl at pH 8.5, 40 mM 2-chloroacetamide (CAA) and 10 mM Tris(2-carboxyethy)phosphine-hydrochloride (TCEP). The Urea buffer contained 8 M Urea, 100 mM Tris-HCl at pH 8.5, 100 mM NaCl, 1mM EDTA, 1X Protease inhibitor cocktail and 1X phosphatase inhibitor PhosphoSTOP. The SDS buffer contained 4% SDS, 100 mM Tris-HCl at pH 8.5, 40 mM CAA and 10 mM TCEP. All the samples were stored at -80°C until further processing.

### EGF stimulation experiments

HeLa cells were grown to ∼70% confluence in DMEM medium supplemented with 10% FBS in 15 cm petri dishes. Cells were stimulated with 50 ng/ml recombinant human epidermal growth factor (Cat No 11343406, ImmunoTools) for 15 and 60 minutes or left untreated, washed in ice cold PBS and lysed in 4% SDC buffer without CAA and TCEP.

### Quizartinib treatment experiments

FLT3-ITD receptor kinase expressing Molm13 and MV4-11 cells were collected at a density of 1 million cells per ml in RPMI medium supplemented with 10%FBS. The cells were washed in plain RPMI medium twice and treated with 20nM quizartinib (gift from Prof. Dr. Frank D Böhmer, Jena) for 4 hours. After 4 hours of incubation, cells were washed twice with ice cold PBS, cell pellets were lysed in 4%SDC buffer, and stored at -80°C until further processing.

### AML patient samples

AML samples were obtained in accordance with the Declaration of Helsinki. All patients were provided informed consents prior to sample collection at the department of hematology, University Hospital Jena and the study was approved by the ethics committee [Vote 3477-06/12]. Briefly, retrospectively the peripheral blood samples collected (during 2012-2016) were enriched for mononuclear cells using density-gradient centrifugation for 30 mins at 400 x g using BIOCOLL separating solution (Density 1.077, Biochrom, Germany). The blast-enriched PBMCs were washed in 1X PBS and preserved at -150 °C. For phosphotyrosine analysis, the cryopreserved samples were thawed as described earlier (Jayavelu et al., 2016). The cells were lysed in 200µl-400µl of SDC lysis buffer (4%SDC,100mM Tris pH8.5, 40mM CAA and 10mM TCEP), BCA quantified and 2mg of samples were processed as described below in the EasyAb sample preparation workflow.

### Animal experiment for GPCR signaling

Female prodynorphin knockout mice (Loacker et al., 2007) (3 mice per treatment; 3 months old) were injected intra-cisternally with either saline, or 20 nmoles of U-50488H, or 30 nmoles of 6’-GNTI in a fixed volume of 3 µl under mild sevofluorane anesthesia. Animals were sacrificed 5 min after injection by cervical dislocation, brains removed and immediately micro-dissected to obtain samples from cortex, hippocampus, striatum, and medulla oblongata. For PP2 inhibitor treatment, animals were injected with PP2 at a dose of 5mg/kg for 30 mins before saline or 6’GNTI. Samples were snap-frozen in liquid nitrogen and stored at – 80°C until further analysis.

### Animal experiment for leptin signaling

Experiments were performed in adult male C57BL/6J mice purchased from Janvier Labs (Saint-Berthevin, Cedex, France). Mice were maintained on a 12-h light-dark cycle with free access to water and standard chow diet (Altromin, #1314). Mice were subjected to a single intraperitoneal (i.p.) injection of either vehicle or leptin, 30 min before being sacrificed by cervical dislocation for organ withdrawal. Brains were extracted swiftly, the hypothalamus was dissected and flash frozen in 40 µl SDC buffer (4% sodium deoxycholate and 100mM Tris pH8.5 without TCEP and CAA). Recombinant murine leptin (R&D Systems, Wiesbaden, Germany) was reconstituted in 20 mM Tris-HCI, pH 8.0 at a concentration of 5 mg/ml. This was then further diluted in saline (0.9% NaCl) to a final concentration of 1 mg/ml and injected at a dose of 3 mg/kg body weight. All studies were approved by the State of Bavaria, Germany.

### Measurement of plasma leptin from mice

Blood from mice was collected in EDTA-containing centrifuge tubes and centrifuged at 4°C and 2000xg for 10 min. Plasma leptin levels were measured with the murine leptin ELISA kit (Merck Millipore, Darmstadt, Germany) according to the manufacturer’s instructions.

### Immunoprecipitation and Western blotting for leptin regulated sites

For detection of leptin-regulated sites, the dissected hypothalami obtained from mice were flash frozen and proteins from the tissue were extracted in freshly prepared RIPA buffer (1% NP40,150 mM NaCl, 50 mM Tris-HCl at pH7.4, 1 mM EDTA, 0.25% deoxycholic acid, 1X phosSTOP, and 1X protease inhibitor cocktail). The lysates were centrifuged at 13,000 rpm g for 10 minutes at 4°C and debris was discarded. For Co-IP, the supernatants were incubated with indicated primary antibody at 4 °C overnight, followed by an incubation with protein G-sepharose beads for 2 hours. The beads were washed in lysis buffer thrice, dissolved with loading buffer and boiled at 95°C for 5 mins. For western blotting, the supernatants were resolved on a 4-12% SDS-PAGE gel and transferred to nitrocellulose membrane. The blots were blocked with 2% BSA-TBST buffer, probed with the indicated primary antibodies (JAK2 (#3230), JAK2 Y1007/1008 (#3771), JNK (#9252), JNK Y185 (#4668), FGFR1 (#9740), FGFR1 Y653/Y654 (#52928), SIRPA (#13379), all purchased from CST), followed by the secondary antibody (HRP conjugated secondary), and developed using enhanced chemiluminescence.

### Cell proliferation assay and drug screen

AML cell lines were seeded in 96well plates at a seeding density of 3 ×10^5^ cells/ml and incubated for 72 hours with increasing concentrations of CDK1/2/5 inhibitor Dinaciclib (S2768), CDK4&6 inhibitor Palbociclib (S1116) and Src inhibitor Bosutinib (S1016, Selleckchem) as illustrated in the figures. Cell growth was assessed after 72 hours by treating with MTS (Cell Titre 96 Aqueous one solution reagent, Promega, Mannheim, Germany) for 4 hours and plates were measured at 490nm for optical density. Values for 8 technical replicates were obtained, corrected by medium only background and normalized to controls. At least two to three independent experiments were performed to assess reproducibility.

### Growth assay

AML cell lines were seeded in 12 well plates at a seeding density of 3×10^5^ cells/ well in appropriate cell culture medium containing 10% heat inactivated FBS and treated with the indicated inhibitors for 48 hours. Viable cells were counted with the trypan blue exclusion method for untreated and inhibitor treated samples. Values for 3 biological replicates were obtained and represented as the mean ± STD.

### Peptide recovery estimation post precipitation and desalting

HeLa cells were grown to ∼70% confluence in DMEM medium supplemented with 10% FBS in 15 cm petri dishes. Cells were collected, washed with ice cold PBS, and lysed in 8M Urea buffer (8M Urea, 100mM Tris pH8.5, 100mM Nacl, 1mM EDTA, 1X Protease inhibitor cocktail and 1X PhosphoSTOP), incubated for 10 minutes on ice, and sonicated for 15minutes at 4°C using Biorupter plus. The sample was centrifuged, and the clear supernatant was carefully transferred to a fresh 15 mL tube. Protein concentration was determined using BCA and Tryptophan fluorescence assay. Next samples were carbamidomethylated using 10 mM TCEP (tris(2-carboxyethyl) phosphine) and 40 mM CAA (Chloroacetamide) for 20mins. Likewise, cells were also lysed in 4%SDC buffer containing 100mM Tris pH 8.5, 40mM TCEP and 10mM CAA, boiled at 95°C for 5 minutes, sonicated for 15 minutes at 4°C using Biorupter plus and processed.

The sample were divided into 1,2,5 and 10 mg protein amounts in 5 mL tubes, diluted to 50% with milliQ water, and precipitated with overnight at -20°C using 4 volumes of Acetone (stored at -20°C). Precipitated proteins were collected by centrifugation at 4000g, 4°C for 10min, pellets washed twice with 80% acetone at -20°C (with a brief sonicate step of 5 minutes to dislodge and disperse the pellets). After centrifuging at 4000rpm for 10mins, the supernatant was discarded and samples dried on paper towel for a few minutes at room temperature until no residual acetone remained. Protein concentration was then determined using BCA and Tryptophan fluorescence assay.

The pellets were resuspended in 20% TFE digestion buffer (20 % 2,2,2, trifluoroethanol, 100mM ammonium bicarbonate, 40mM CAA and 10 mM TCEP) with sonication (Biorupter for 10minutes or a until homogenous suspension was formed). Again, protein concentration was determined using BCA and Tryptophan fluorescence assay. Proteins amounts 1,2,5 and 10 mg were digested with trypsin (1:100 ratio, Sigma) and LysC (1:100 ratio, Sigma) at 37°C overnight. The digested peptides were desalted using the Sep-Pak Vac-1cc 100mg and Sep-Pak Vac-6cc 500mg Sorbent per tC18 Cartridge (Waters, Milford, MA, USA). Column-conditioning, peptide desalting and elution steps were performed step wise as described below.

### Desalting of digested peptides

Prior to desalting, peptide quantity was measured using the tryptophan fluorescence assay. Peptides were acidified with 0.5% TFA. Cartridges were first conditioned with 80% acetonitrile and 0.5% acetic acid, then equilibrated with 0.5% acetic acid. The acidified peptides were loaded onto the cartridges, washed with 0.2% TFA, and eluted with 80% acetonitrile containing 0.5% acetic acid (centrifuged at 50 xg for 1 minute per step). Samples were dried overnight in a speed vacuum. The dried peptides were reconstituted in 2% acetonitrile and 0.1% TFA, and peptide concentration was verified by tryptophan fluorescence. Purification efficiency for protein precipitation by acetone and peptide desalting was determined based on measured protein and peptide yields.

### Immuoprecipitation and western blotting for EasyAb workflow development

Protein extraction and digestion was carried out by lysing the cells in 4%SDC buffer (4%SDC and 100mM Tris pH 8.5 without TCEP and CAA). The lysates were sonicated for 10min using Biorupter plus (Diagenode) followed by 10 min of incubation on ice. Protein concentration was determined by BCA and subsequently the samples were alkylated with 40 mM CAA and 10mM TCEP in the dark for 30minutes on ice. For direct immunoprecipitation of proteins, the concentration of SDC buffer was adjusted to 0.4% by diluting the 4%SDC buffer with milliQ water. For immunoprecipitation experiments: the proteins were digested with trypsin (1:100 ratio, Sigma) and LysC (1:100 ratio, Sigma) for 4 hours at 37°C. This was followed by antibody-based peptide enrichment for phosphotyrosine peptides using anti-phosphotyrosine pY1000 antibody conjugated to magnetic beads (Cat # 14017, Cell Signaling Technology). The enriched peptides were boiled in sample buffer (NuPAGE, Invitrogen). The samples were then resolved on a 4-12% PAGE gel, and stained with either coomassie or silver stained or alternatively blotted on PVDF membrane, blocked with 2%BSA TBST buffer and developed using LAS (GE).

For desalting prior to post translationally modified peptide enrichment, Sep-Pak Vac-6cc 500 mg Sorbent per tC18 cartridges with 37-55 µm particle size (WAT036790, Waters, Milford, MA, USA) were used. The cartridge conditioning was performed step wise and twice with 3 ml of methanol, 3 ml of 80% acetonitrile (ACN) and 3 ml of 0.2% trifluoro acetic acid (TFA). The columns were then loaded with trypsin/LysC-digested peptides for desalting, washed with 0.2% TFA twice and eluted with 3 ml of 100% and then 3 ml 80% ACN. The peptides were vaccum-dried (using a speedvac) and resuspended in IAP buffer (50 mM Tris-HCl at pH7.4, 150 mM NaCl, 1% NOG (n/octyl-ß-D-glucopyranoside) supplemented with 1X protease inhibitor and 1X phosphatase inhibitors. PTM peptide enrichment was carried out by incubating the desalted peptides with anti-phosphotyrosine pY1000 antibody.

### Tissue and cell lysis in conventional buffer

HeLa cells lysed in 4%SDS buffer (4%SDC, 100mM Tris pH 8.5 , 40mM TCEP and 10mM CAA). were heated for 5 minutes at 95°C and stored at -80°C until further processing. The lysates were sonicated for 15minutes on Biorupter plus (Output-High, 30 second on/off @4°C) and heated again at 95°C, followed by a brief 1-minute centrifugation at 13,000 rpm. For sample preparation in Urea buffer: HeLa cells were lysed in 8M Urea buffer (8M Urea, 100mM Tris pH8.5, 100mM Nacl, 1mM EDTA, 1X Protease inhibitor cocktail and 1X PhosphoSTOP), incubated for 10 minutes on ice, and sonicated for 15minutes at 4°C using Biorupter plus. Samples were then incubated with 40mM CAA for 30minutes in dark, followed by 10mM TCEP for 15mins on ice.

The supernatants were removed into a clean tube, diluted to 50% with milliQ water, and precipitated with overnight at -20°C using 4 volumes of Acetone (stored at -20°C). Precipitated proteins were collected by centrifugation at 4000g, 4°C for 10min, pellets washed twice with 80% acetone at -20°C (with a brief sonicate step of 5 minutes to dislodge and disperse the pellets ). After centrifuging at 4000rpm for 10mins, the supernatant was discarded and samples dried on paper towel for a few minutes at room temperature until no residual acetone remained. The pellets were resuspended in 20% TFE digestion buffer (20 % 2,2,2, trifluoroethanol, 100mM ammonium bicarbonate, 40mM CAA and 10 mM TCEP) with sonication (Biorupter for 10minutes or a until homogenous suspension was formed). Protein concentration was determined using BCA.

### Protein digestion, desalting, and PTM peptide enrichment for conventional workflows

Proteins amounts as described in the text, figures or in the legends were digested with trypsin (1:100 ratio, Sigma) and LysC (1:100 ratio, Sigma) at 37°C overnight. The digested peptides were desalted using the Sep-Pak Vac-6cc 500mg Sorbent per tC18 Cartridge. Column-conditioning, peptide desalting and elution steps were performed step wise as described above except that the peptides were eluted with only 100% acetonitrile. The desalted peptides were vaccum-dried and subsequently resuspended by agitation in ice cold IAP buffer (with1X phosphoSTOP and 1X protease inhibitor cocktail) for one hour.

Posttranslationally modified peptides of interest were enriched using antibodies against phosphotyrosine antibody (anti-phosphotyrosine pY1000 antibody conjugated to magnetic beads (100 µl/sample)), anti-acetylated-lysine antibody (5 µl of Ac-K2-100 MultiMab ^TM^ Rabbit mAb mix /1 mg peptide) or anti-monomethylated arginine antibody (5 µl of mme-R MultiMab^TM^ Rabbit mAb mix /1 mg peptide) Incubation of the peptides with the antibodies occurred at 4°C overnight with rotation. For unconjugated antibodies, pre-washed protein A/G magnetic bead slurry was added (20 µl/ mg peptide, Cat No 88802, Pierce) and incubated on rotation for 2 hours at 4°C. The bead-bound PTM-peptides were washed three times with ice cold IAP buffer and three times in milliQ water by pelleting the beads using the magnetic separation rack (DynaMag-2, ThermoFisher Scientific).

To release the antibody bound modified peptides, 200µl of 0.2% TFA was added and incubated for 10-15 minutes at room temperature. Peptides were collected using a magnetic rack and the procedure was repeated twice. Peptides were then loaded onto equilibrated (30% methanol, 1% TFA) styrenedivinylbenzene-reversed phase sulfonated (SDB-RPS) Stage Tips (Rappsilber et al., 2003) for desalting. Peptides were subsequently washed with 0.2% TFA and eluted into clean PCR tubes with 60 µl of elution buffer (80% of Acetonitrile, 1.25% NH4OH). The eluates were vaccum-dried, resuspended in MS loading buffer (3% ACN, 0.3% TFA) and stored at -20°C until MS measurement.

### EasyAb sample preparation workflow

Cells or tissues equivalent to 2 mg or more were lysed in 200 µl 4% SDC lysis buffer (4% SDC, 100mM Tris pH8.5 without TCEP and CAA), heated for 5 minutes at 95°C and stored at -80°C until further processing. The samples were sonicated for 15 min using Biorupter plus followed by protein concentration determination using BCA. (NOTE: TCEP might interfere in the BCA reaction, therefore the reduction step must be carried out after protein concentration estimation). Subsequently, the samples were alkylated with 40 mM CAA and 10 mM TCEP in the dark for 30 min on ice.

Two milligrams of proteins (unless indicated otherwise) were digested with trypsin (1:100 ratio, Sigma) and LysC (1:100 ratio, Sigma) for 4 to 12 hours at 37°C. The digested peptides were diluted to 4x the initial volume with double distilled water and another 4x with IAP buffer containing 1X PIC (protease inhibitor cocktail) (NOTE: addition of 1X PIC aids to inactivate any residual protease activity of Trypsin and LysC), followed by pH adjustment to 7.4 using 1 M Tris-HCl buffer (pH 7.4) (NOTE: typically 50 µl of 1 M Tris-HCl at pH7.4 is sufficient for 1.8 ml of lysate).

Posttranslationally modified peptides of interest were enriched using the antibodies against phosphotyrosine antibody (anti-phosphotyrosine pY1000 antibody conjugated to magnetic beads (100 µl/sample)), anti-acteylated-lysine antibody (5 µl of Ac-K2-100 MultiMab ^TM^ Rabbit mAb mix /1 mg peptide) or anti-monomethylated arginine antibody (5 µl of mme-R MultiMab^TM^ Rabbit mAb mix /1 mg peptide Incubation of the peptides with the antibodies occurred at 4 hours (at room temperature) or overnight (at 4°C) while rotating. For unconjugated antibodies, pre-washed protein A/G magnetic bead slurry (Cat No 16-663, Merck) was added ( 20µl / mg peptide) and incubated on rotation for 2 hours at 4°C. The bead-bound PTM-peptides were washed three times with ice cold IAP buffer and three times in milliQ water by pelleting the beads using the magnetic separation rack.

To release the antibody bound modified peptides, 200µl of 0.2% TFA was added and incubated for 10-15 minutes at room temperature. Peptides were collected using a magnetic rack and the procedure was repeated twice. Peptides were then loaded onto equilibrated (30% methanol, 1% TFA) styrenedivinylbenzene-reversed phase sulfonated (SDB-RPS) StageTips for desalting. Peptides were subsequently washed with 0.2% TFA and eluted into clean PCR tubes with 60 µl of elution buffer (80% of acetonitrile, 1.25% NH4OH). The eluates were vaccum-dried, resuspended in MS loading buffer (3% ACN, 0.3% TFA) and stored at -20°C until MS measurement.

### High throughput EasyAb workflow

Digested peptides (2 mg in 200µl) were transferred to a clean 2 ml 96-well deep-well plate (DWP with white border, Eppendorf, Cat No 0030504305) and the SDC concentration was adjusted to 0.4% by diluting the samples with 800 µl of double distilled water (4x volume) and 800 µl of IAP buffer (4x volume), followed by a pH adjustment to 7.4. To perform phosphotyrosine peptide enrichment, 100 µl of magnetic beads conjugated anti-phosphotyrosine pY1000 antibody was added as mentioned previously. The DWP was sealed with heat-sealing film (Eppendorf, Cat No 0030127838) using PCR system Heat Sealer S200 (Eppendorf) at 200°C for 2 s and placed for rotation at room temperature for 4 hours. The bead bound peptides were washed three times in IAP buffer and twice in double distilled water by placing the DWP on a Magnum EX Universal Magnet Plate (Alpaqua, Cat No SKU A000380). The beads were collected in 200 µl double distilled water and transferred to a 96 well PCR plate (Eppendorf twin.tec PCR plates 96 LoBind, skirted), followed by removing the water by placing the plate on 96 side skirted magnet (DynaMag, Cat No 12027). The beads-bound peptides were released by adding 200µl of 0.2% TFA to the PCR plates and incubated for 10-15 minutes at room temperature. The released peptides were collected using the 96 Side skirted magnet. Further desalting on SDB-RPS StageTips, elution and storage of peptides were done as mentioned above.

### Single run LC-MS/MS measurements

All samples were analyzed on nanoflow HPLC coupled to a high resolution mass spectrometer. In brief, the peptides were loaded onto a 50-cm column with 75-µM diameter, packed in house with 1.9 µM C18 ReproSil particles (Dr. Mischa GmbH). The column temperature was maintained using a homemade column oven at 50°C. An EASY-nLC 1000 system (Thermo Fischer Scientific) was connected to the mass spectrometer with a nano spray ion source, and the peptides were separated with the binary buffer system of 0.1% formic acid (buffer A) and 60% ACN plus 0.1%formic acid (buffer B), at a flow rate of 300nl/min. Peptides were eluted over 140 minutes with a gradient of 5-20% buffer B over 85 minutes, 20-65% over 45minutes, and 65-80% over 10 minutes. Samples were analyzed on a Q Exactive HF-X Hybrid quadrupole-Orbitrap mass spectrometer (Thermo Fischer Scientific), with one full scan at a target of 3e^6^ ions (300-1650 m/z, R=60,000 at 200 m/z) , followed by Top10 MS/MS scans with HCD (high energy collisional dissociation) (target 1e5 ions, maximum filling time 120ms, isolation window 1.6 m/z, and normalized collision energy 27%) and detected in the Orbitrap (R=15,000 at 200 m/z). Dynamic exclusion for 40s, Apex trigger 4s to 7s and charge exclusion (unassigned, 1,5, -8 & >8) were enabled.

### Deep proteome measurements

Cell or tissues (20µg in 50µl) were lysed in 1% SDC buffer (1% SDC, 100mM Tris pH8.5, 40mM CAA, and 10mM TCEP) on ice for 20 min, heated at 95°C for 5 min, sonicated for 15 minutes, and heated again at 95°C for 5 minutes. Proteins were digested with trypsin (1:100 ratios, Sigma) and LysC (1:100 ratios, Sigma) at 37°C for 4 hours. To stop the digestion, 5x volume isopropanol/1% TFA was added and vortexed vigorously. The peptides were desalted on SDB-RPS StageTips, washed with 0.2% TFA and eluted with 60µl of elution buffer (80%, 1.25% NH4OH). The dried eluates were resuspended in MS loading buffer (3%ACN, 0.3% TFA) and stored at -20°C until MS measurement.

### Data analysis

All raw files were processed using Maxquant (Cox and Mann, 2008) version 1.5.5.2 supported by the Andromeda search engine. The data was searched for proteins and peptides using a target-decoy approach with a reverse database against Human Uniprot (2016) or Mouse Uniprot (2016) fasta file with a false discovery rate of 1% at the levels of protein, peptide and modification. We used default settings with the following minor changes: oxidized methionine (M), acetylation (protein N-term) and in case of phosphopetide search: phospho (STY), for mono methyl arginine: mono methyl on lysine and arginine, for acetylated lysine: acetyl lysine were selected as variable modifications, and carbamidomethyl (C) as fixed modification. A maximum of 2 missed cleavages were allowed, and a minimum peptide length of seven amino acids was set. The peptides were identified with an initial precursor mass tolerance of up to 7ppm and fragment mass tolerance of 20ppm. Match between run algorithm was enabled were ever applicable (requires minimum 3 samples per condition or group) with a matching window of 1 min. For label free protein quantification the minimum ratio count was set to 2.

### Bioinformatic analysis and statistics

MaxQuant output tables were processed with Perseus (Tyanova et al., 2016) version 1.5.2.11, InstantClue (Nolte et al., 2018), and GraphPad Prism 7 for statistical and clustering analysis. The Annotations were extracted from Gene Ontology (GO), Uniprot-Keywords, Kyoto Encyclopedia of Genes and Genome (KEGG), Phosphosite plus, Wikipathway and Reactome. Protein interaction network of experimentally validated EGF regulated phosphoproteins were analyzed using the STRING interaction database and visualized in Cytoscape (Szklarczyk et al., 2019). The study consistently utilized biological replicate samples for reproducibility and no samples were removed from the analysis. Significance was assessed by Student’s t-test, using permutation –based FDR to control for multiple hypothesis testing. The p-values were calculated using a paired two tail Student’s t-test.

### Data

All the MS raw files were deposited to the ProteomeXchange Consortium via the PRIDE partner repository and available with the identifier PXD015503. The reviewer account details are as follows

