## Supplemental Information for "EasyAb: A High-Throughput Workflow for Antibody-Based PTM Peptide Enrichment Method Coupled to Mass Spectrometry"

**SUPLEMENTARY TABLES**

**Table S1.** Benchmarking EasyAb vs Conventional Workflow

**Table S2.** AML patient characteristics

**Table S3.** AML patient FLT3WT vs FLT3ITD.xls

**Table S4.** Leptin regulated sites in hypothalamus.xls

### SUPPLEMENTARY

#### Establishing the EasyAb workflow.

Having successfully benchmarked the EasyAb-SDC buffer system for PTM peptide enrichment, we further wanted to test whether desalting the SDC buffer prior to enrichment provides any advantage in the peptide quantification. To this end we compared direct SDC lysis and enrichment (from 2 mg lysates) procedure to desalting the peptides after protease digestion in SDC buffer for the enrichment (from 2 mg peptide) of phosphotyrosine modified peptides from MV4-11 and Molm13 cells. In addition we also tested whether antibody amounts affect the enrichment & quantification (**Figure S2A**). In brief, there was no significant difference in number of quantified peptides between direct and desalting procedure (**Figure S2D,E**) and this was supported by high reproducibilty between the samples (**Figure S2F**).

#### Visualizing FLT3 ITD phosphotyrosine signaling in AML cells

With the deep phosphotyrosine phosphoproteome obtained from human MV4-11 and Molm13 cells we here visualized the FLT3-ITD mutant tyrosine kinase receptor signaling (**Figure S3**).

FLT3 belongs to the class-III type tyrosine kinase receptor family, plays an important role in differentitaion, proliferation and survival of hematopoietic stem and progenitor cells and thereby tightly regulates hematopoiesis. The Internal Tandem Duplication (ITD) mutation in the FLT3 receptor leads to its ligand independent activation, and this mutations occurs in approximately 20% of AML. Notably, patients with FLT3-ITD respond very poorly to standard chemotherapy. Therefore understanding the signaling and cross-talk relayed by FLT3-ITD in AML cells is therapeutically essential to target it. We quantified five phosphotyrosine sites in FLT3 receptor (Y401, Y630, Y726, Y842 & Y969), among these sites Y401 is novel, whereas the other four sites are already known to have important regultory function. Autophosphorylation of Y726 in the kinase catalytic domain is required for activation and phosphorylation of FLT3 substares, phosphorlation on Y842 is required for activation of the kinase and promotes growth and anti-apoptotic signals whereas phosphorlation on Y969 regultes the cell cycle (Choudhary et al., 2005; Kindler et al., 2005; Razumovskaya et al., 2009; Schmidt-Arras et al., 2005).

Constitutively active FLT3 recruits and tyrosine phosphorylates adaptor proteins to signal to its downstream targets. We found several adaptor proteins phosphorylation such as Gab1 (Y659), Gab2 (Y293,Y476,Y614, Y643), Gab3(Y560), Grb2 (Y209), Inddp5 (Y1161), Nck1 (Y105, Y112), Nck2 (Y105), Pi3kap1 (Y570, Y694) and Shc1(Y427). These tyrosine phosphorylated adaptor proteins further recruit and transduce FLT3-ITD mediated signaling cascades via the SH2 and SH3 docking domains they encompass. In this manner, FLT3 activates Ras-Mapk pathway members for aberrant cellular proliferative and survival signals via recruitment and activation of adaptor proteins Grb2 and Shc1(Marchetto et al., 1999; Mizuki et al., 2000). For PI3-K pathway activation, FLT3 was reported to utilize Gab1, Gab2 or Ptpn11 (Zhang and Broxmeyer, 1999). These adaptor are thought to be the key for Pi3k activation since human FLT3 lacks the analogous PI3K consensus site required for directly Pi3K interaction.

Activation of Stat3 and Stat5 was shown to be critical for FLT3-ITD mediated leukemia transformation (Rocnik et al., 2006). We quantified Stat3 (Y705) and Stat5 (Y694) phosphorylation, in addition we observed STAT6 Y461 phosphorylation in FLT3-ITD cells. The biological consequence of Stat6 signaling is the activation of innate immunity (Takeda et al., 1996). We recently showed that Stat6 is a direct substrate of Ptpn6 in FLT3-ITD cells and that Ptpn6 depletion promotes Stat6 activation (Reich et al., 2020). Loss of STAT6 in FLT3-ITD cells accelerates leukemia like disease in vivo. However, interestingly, it is not yet known which upstream kinase phosphorylates STAT6 and what type of innate immune signatures are triggered in FLT3-ITD cells.

Several tyrosine kinases such as Syk , Btk, Src kinase family (SFKs) activation have been shown to be critical for FLT3-ITD depended disease progression. We quantified nearly 30 tyrosine kinases activation sites, many of which were reported to be therapeutically important and involved in cancer drug resistance but not previosuly known to be regulated by FLT3-ITD. TNK2 also known as ACK1, is an oncokinase whose activity was linked to tamoxifen-resistance breat cancer and castration-resistance prostate cancer (Mahajan et al., 2017; Wu et al., 2017). We quantified multiple phosphorylation sites on TNK2 (Y40, Y284,Y827,Y857,Y859), of which Y284 is required for activation. A known target of TNK2, Histone H4 phosphorylation at Y88 was also found to be phosphorylated in FLT3-ITD cells (Mahajan et al., 2017). This provides evidence that TNK2 kinase mediated Histone H4 Y88 marks might play a role in FLT3-ITD signaling.

Receptor tyrosine kinase Axl was previously shown to be consitutively active in AML cells and high levels of Axl expression are assocaited with worse survival (Rochlitz et al., 1999). Activated Axl potentially acts as a docking site for Syk and Pi3k in leukemic cells (Gioia et al., 2015). Interestingly, Axl was shown to be critcial for response to tyrosine kinase inhibitor PKC412 in AML cells with FLT3-ITD (Park et al., 2015). We quantified Axl (Y702, Y703, Y866) tyrosine phosphoryaltion. However the relevance of these tyrosine phosphorylation is still unclear. Conversely, our data now supports the notion that FLT3-ITD might establish the cross-talk with Axl that benefits AML cells for TKi mediated drug response and survival.

Reactive Oxygen Species (ROS) generation is one of the critical components for FLT3-ITD mediated transformation (Jayavelu et al., 2016). Catalase helps in neutralizing reactive hydrogen peroxide, the primary ROS produced in FLT3-ITD cells. We quantified phosphorylation on Catalase at several tyrosine sites

(Y84,Y187, Y231, Y308, Y379), of which Y231 is required for induction of enzyme activity (Cao et al., 2003), indicating  FLT3-ITD directly regulates Catalase enyzme activity through Y231 phosphorylation in AML cells.

**AML cell line specific phosphotyrosine signature and sensitivity to Dinaciclib (CDK1/2/5/9 inhibitor).**

Among the quantified phosphotyrosine sites in patient samples and cell lines (**Figure S5F,G**), we focused on the driver gene mutation specific changes in cell lines (since these cell lines have publically available targeted sequencing data from Broad Institute Cancer Cell Line Encyclopedia). Notably, we have captured the known tyrosine phosphorylation of the driver kinases in the corresponding cell lines such as Pdgfr (Y894) in EOL1 cells, Jak2 (Y570) in HEL cells, c-Kit (Y609) in Kasumi-1 cells, Abl (Y226, Y264, Y393) in SD1 cells (**Figure S7A**). Further we quantified mutation specific changes that had not previously been reported such as Rap2c Y166 in HL60 cells, Cdk Y15 in ML2 cells, Fam83H Y1032, Ncor2 Y1270 in OCI-AML3 cells and Lilrb 4 Y442 in OCI- AML5 cells (**Figure S7A**). To investigate whether these phosphorylation changes are therapeutically relevant, we chose to validate the druggable kinase action of CDK. Deregulated CDK activation results in abnormal cell cycle progression and small molecules targeting CDK show anti-tumor effects in various cancer models. Phosphorylation of CDK1,2,3 and5 on Y15 leads to inhibition of enzyme activity and constitutes an anti-apoptotic signal. We observed varying degrees of CDK1/2/3 Y15 phosphorylation between the cell lines we screened but no phosphorylation on CDK4 and 6. Therefore to test whether the cell lines are CDK1/2/3 dependent we treated then with increasing concentration of the CDK1,2,5&9 inhibitor Dinaciclib and the CDK4&6 inhibitor Palbociclib. The growth assay revealed a selective vulnerability and high sensitivity to Dinaciclib but not to Palbociclib (**Figure S7B, C**). Moreover, we found ML2 cells to have greater sensitivity to Dinaciclib (IC50 at ~5 nM) than other tested cell lines. This data shows that the response to Dinaciclib is more dependent on the degree of Y15 phosphorylation than that of CDK protein expression between the corresponding cell lines (**Figure S7E**). We further confirmed this notion by measuring the rate of apoptosis in the cell lines and found ~90% cell death in ML2 cells compared to 55-75% other cell lines at 8nM Dinaciclib (**Figure S7D**). Of note, Dinaciclib showed 50% kinase inhibitor concentration at nM ranges. Sensitivity to MLL rearranged acute myeloid leukemia was demonstrated previously however it was not tested against non MLL rearranged AML cell lines (Baker et al., 2016). Thus our results suggest the intriguing possibility that Y15 phosphorylated CDK1/2/3 could serve as a marker for Dinaciclib application and that the phosphotyrosine changes we present here could be of potential drug targets/markers for the treatment of AML.

**GPCR signaling**

In the brain, GPCRs, such as kappa opioid receptor (KOR), mediate neurotransmitter metabotropic signaling that is essential in neuronal plasticity and function. KOR is primarily expressed in cortical and sub-cortical regions such as striatum, hippocampus and amygdala. Recently, we applied large-scale $TiO_2$-based phosphoproteomic experiments to discern KOR-mediated signaling pathways that lead to side effects (Liu et al., 2018). From these experiments we explored the brain phospho- serine and threonine architectures in the brain and linked the unwanted side effects to activation of mechanistic target of rapamycin (mTOR). One interesting aspect of those experiments was that activation of tyrosine kinases is also associated with wanted side effects.

Consequently, we applied EasyAb to investigate large scale KOR-mediated tyrosine phosphoproteomic changes from four different mouse brain regions (cortex, hippocampus, striatum, and medulla oblongata) 5 minute after administration of the saline control, U50,488H the aversive agonist, and 6'GNTI the non-aversive agonist. Biological replicates from each brain region were dissected from a single animal (three per treatment). In total, we quantified 350 phospho sites, of which 40% were previously unknown. The number is significantly lower than the cancer cells highlighted above, indicating that tyrosine kinases are more tightly regulated in healthy tissues than their serine and threonine counterparts.

In addition to tyrosine and serine/threonine kinases mentioned in the main text, we also observed tyrosine phosphorylation of Sirpa (Y505) and Tns (Y483) was regulated differently by U50, 488H and 6'GNTI. Furthermore, we found enrichment of Jak2 motifs, indicating that Jak2 may be another signaling component that distinguishes 6'GNTI-induced and U50,488H-induced KOR activation.

Lastly, we observed 6'GNTI specific perturbation of the Src family tyrosine kinase Fyn (pY184, pY213), postsynaptic density protein 95 (PSD95) (pY580, pY609) and components of NMDA or AMPA receptors (**Figure S8B**). Most of these phosphorylation sites bear the Src kinase motif. This could indicate that Src activation may be a player in the communication between KOR activation and glutamate signaling.

**Leptin phosphoproteome signaling**

Apart from the direct observation of Jak2 Y507 autophosphorylation site and the STAT5b Y699 activation site in the dataset, we further found enrichment of Jak2 and Src kinase substrate motifs (**Figure S8F**).

We further found leptin-induced Y2430 phosphorylation of spectrin alpha 1 (Sptan1), a cytoskeletal protein involved in $Ca^{2+}$-calmodulin-regulated vesicular secretion, and Y569 phosphorylation of Ankyrin-2, a protein involved in assembling and stabilizing membrane-bound ion channels and transporters such as Na/Ca exchangers, Na/K ATPases or the Ca2+ channel Inositol 1,4,5 triphosphate receptors (InsP3R). Notably, Ank2-Y569 is a potential Jak2 substrate motif and spectrin family members were shown to interact with InsP3R, potentially indicating a direct role for leptin signaling in regulating membrane transporter and vesicle activity.

Lastly, we identified leptin-induced changes in gene transcription regulators such as Stat5 (pY699), Hipk2 (pY361), ribonucloproetin hnRNPD0 (pY244) and Arhgap35 (pY1105). Surprisingly, we also found significantly regulation of DNA binding protein Ikzf4 on pS22. Ikzf4 plays a role in the development of the central and peripheral nervous system and function as transcriptional repressor, and has not yet been reported as leptin target.

Figure S1

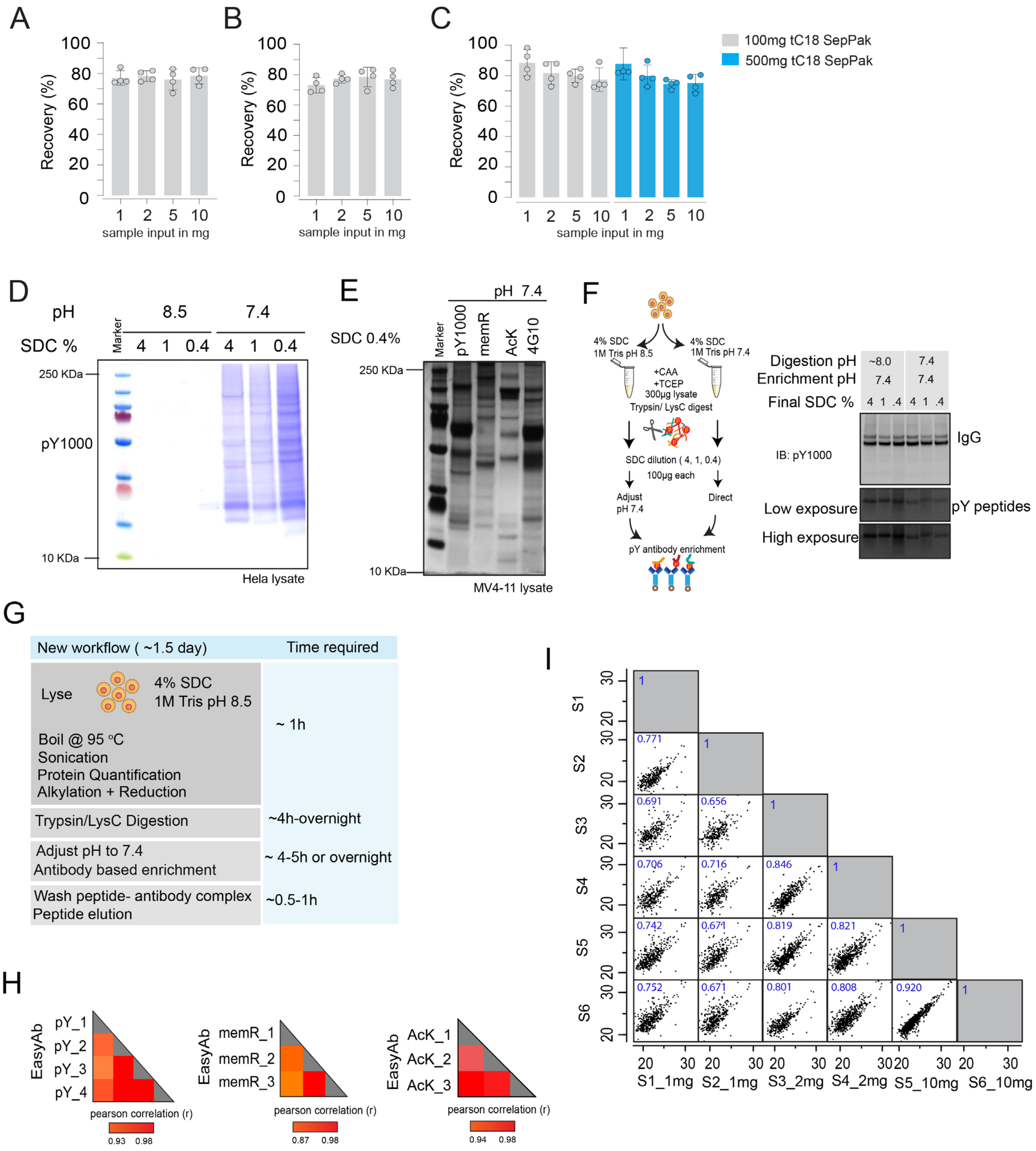


**Figure S1 Development and Optimization of EasyAb workflow.**

(**A**) Protein recovery post acetone precipitation from Hela lysates prepared in 8M Urea. (**B**) Protein recovery post acetone precipitation from Hela lysates prepared in 4%SDS. (**C**) Peptide recovery after desalting using 100 mg and 500 mg C18 desalting cartridges.

(**D**) Optimization of SDC buffer concentration and pH for antibody-based pull down of post translationally modified proteins in HeLa lysate. Coomassie stained SDS-PAGE gel shows enrichment with phosphotyrosine antibody at pH 8.5 and pH 7.4. at different SDC concentrations.
(**E**) Optimum pH and concentration of SDC in EasyAb lysis buffer enables detection of posttranslationally modified proteins in MV4-11 cells. Silver stained SDS-PAGE gel indicates presence of phosphotyrosine-phosphorylated proteins, monomethyl arginine modified proteins, and lysine acetylated proteins immunoprecipitated from MV4-11 cells.
(**F**) Comparison of SDC buffer compatibility for optimal protease digestion and modified peptide enrichment. Left panel: schematic illustration of the protocol. Right panel: 4-12% SDS-PAGE resolved and western blot detected phosphotyrosine peptides (pY) from the lysate when digested using 0.4-4% SDC buffer at pH 8, followed by modified peptides enrichment at pH 7.4. The pY peptides were detected below 10KDa range.
(**G**) Overview of time required to perform EasyAb protocol.
(**H**) Heatmap of Pearson correlation coefficients shows the reproducibility between the samples for pY1000, AcK and memR antibodies using EasyAb workflow in **Figure1C**.
(**I**) Pearson correlation coefficients shows the reproducibility between the samples S1-S6 shown in **Figure 1D** using EasyAb workflow.

Figure S2

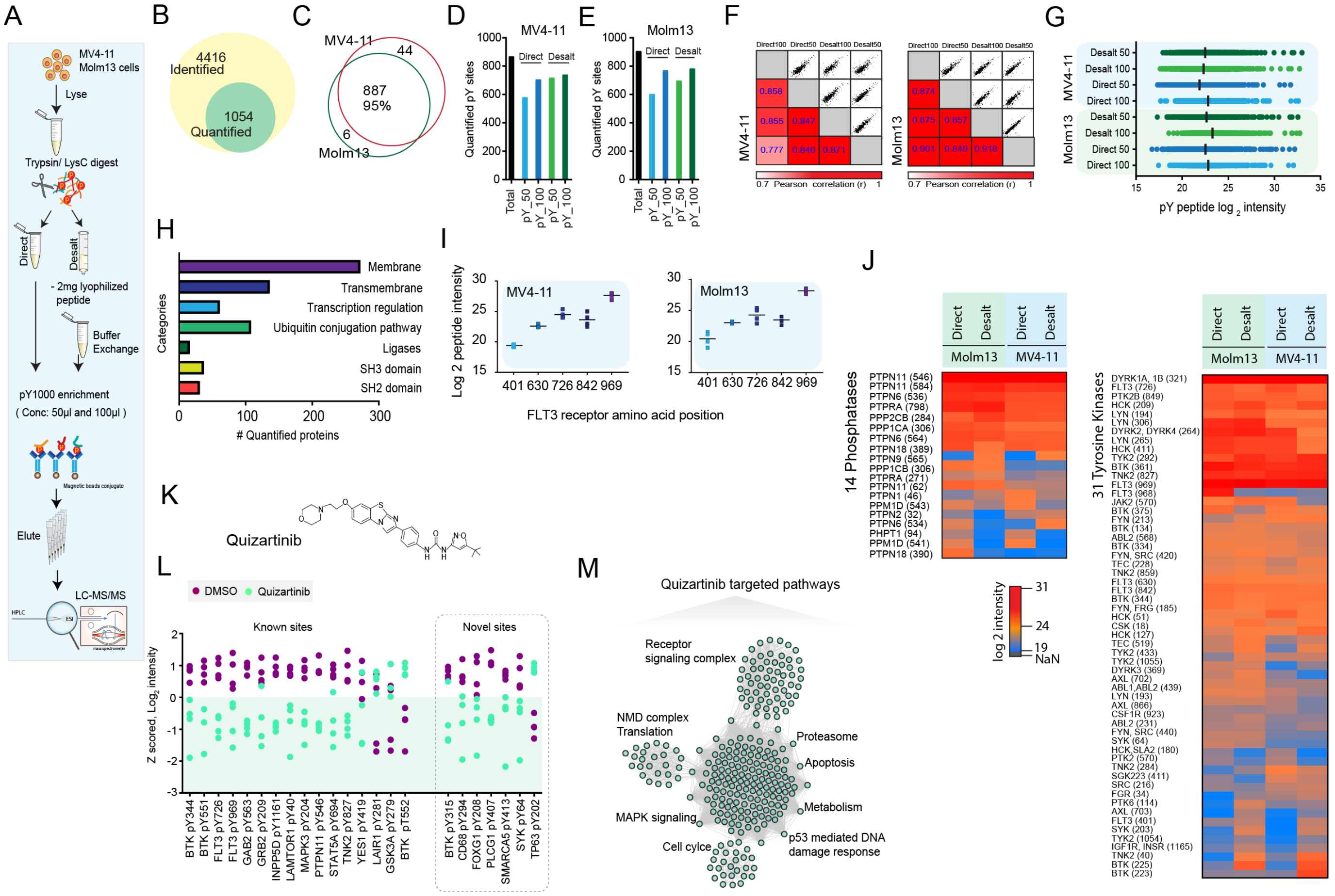


**Figure S2 EasyAb for in-depth phosphotyrosine profiling.**

(**A**) EasyAb workflow scheme. Direct modified peptide enrichment compared to desalting procedure with different antibody amount in MV4-11 and Molm13 cell lines both harboring common mutation in the FLT3 receptor (FLT3-ITD) under their native condition (i.e. without growth factors or sodium pervanadate pretreatment).

(**B**) Total number of identified and quantified phosphosites in MV4-11 and Molm13 cell lines from 2 mg of lysate.

(**C**) The Venn diagram displays the exclusive and shared number of quantified phosphoTyrosine (pY) sites between the MV4-11 and Molm13 cell lines. The phosphotyrosine phosphorylated proteins were 95% common in both the cell lines than the cell line-specific changes highlighting dominance of mutated FLT3 receptor driven tyrosine signaling and success of EasyAb workflow.

(**D-E**) The bar diagram displays the overall number of quantified phosphosites and number of quantified phosphosites in MV4-11 (D) and Molm13 (E) cell lines by comparing direct and desalting procedure under different antibody (magnetic beads conjugated-pY1000) concentrations/volume. This data indicate antibody volume limits the depth of quantification.

(**F**) Heatmap of multiscatter plot, and Pearson correlation coefficients showing the reproducibility between the samples obtained from MV4-11 cell lines.

(**G**) The $log_2$ intensity of the phosphotyrosine sites quantified between the samples and from two cell lines.

(**H**) Bar diagram displays uniprot keyword categories with number of tyrosine phosphorylated proteins. Membrane and Transmembrane proteins were highly enriched in the dataset.
(**I**) Dot plot shows the quantification of FLT3 receptor phosphotyrosine sites. FLT3 Y401 is a novel site quantified in this study.
(**J**) Heatmap shows the $log_2$ phosphosite intensity of quantified tyrosine kinases and phosphatases in the two cell lines.
(**K**) Chemical structure of FLT3 inhibitor Quizartinib.
(**L**) Aligned dot blot shows selected known and novel phosphosites regulated by Quizartinib treatment. The novel sites are highlighted in dotted rectangle box.
(**M**) Network map of significantly enriched GO terms of Quizartinib regulated phosphoproteins in FLT3-ITD cells. Each node represents one GO term and the highlighted terms are most significantly regulated. Quizartinib extensively targets phosphotyrosine phosphorylated proteins annotated for Receptor, MAPK, Cell cycle and Apoptosis signaling.

Figure S3

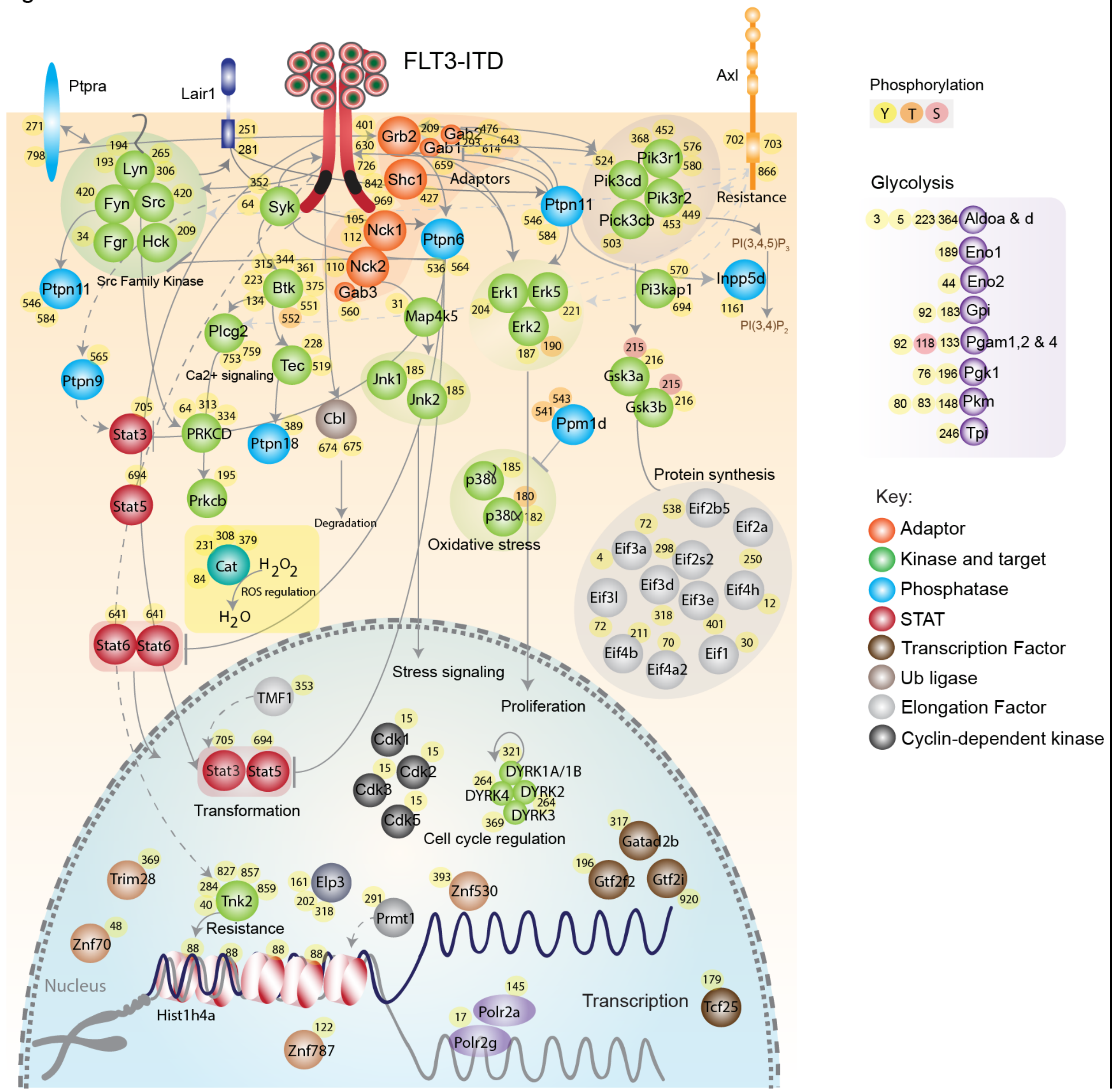


**Figure S3 Visualizing FLT3-ITD mediated signaling in AML cells.**

Illustration of FLT3-ITD mediated phosphotyrosine signaling in the AML cells. The map represents known and novel targets of FLT3-ITD. Majority of the proteins in the network were organized based on their function. Circle on the protein denotes the phosphorylated residue (Tyrosine (Y), Threonine (T) and Serine (S) are represented in different colour as illustrated in the map). Established associations are connected in solid lines/arrows and curated associations are connected in dotted lines/arrows.

FigureS4

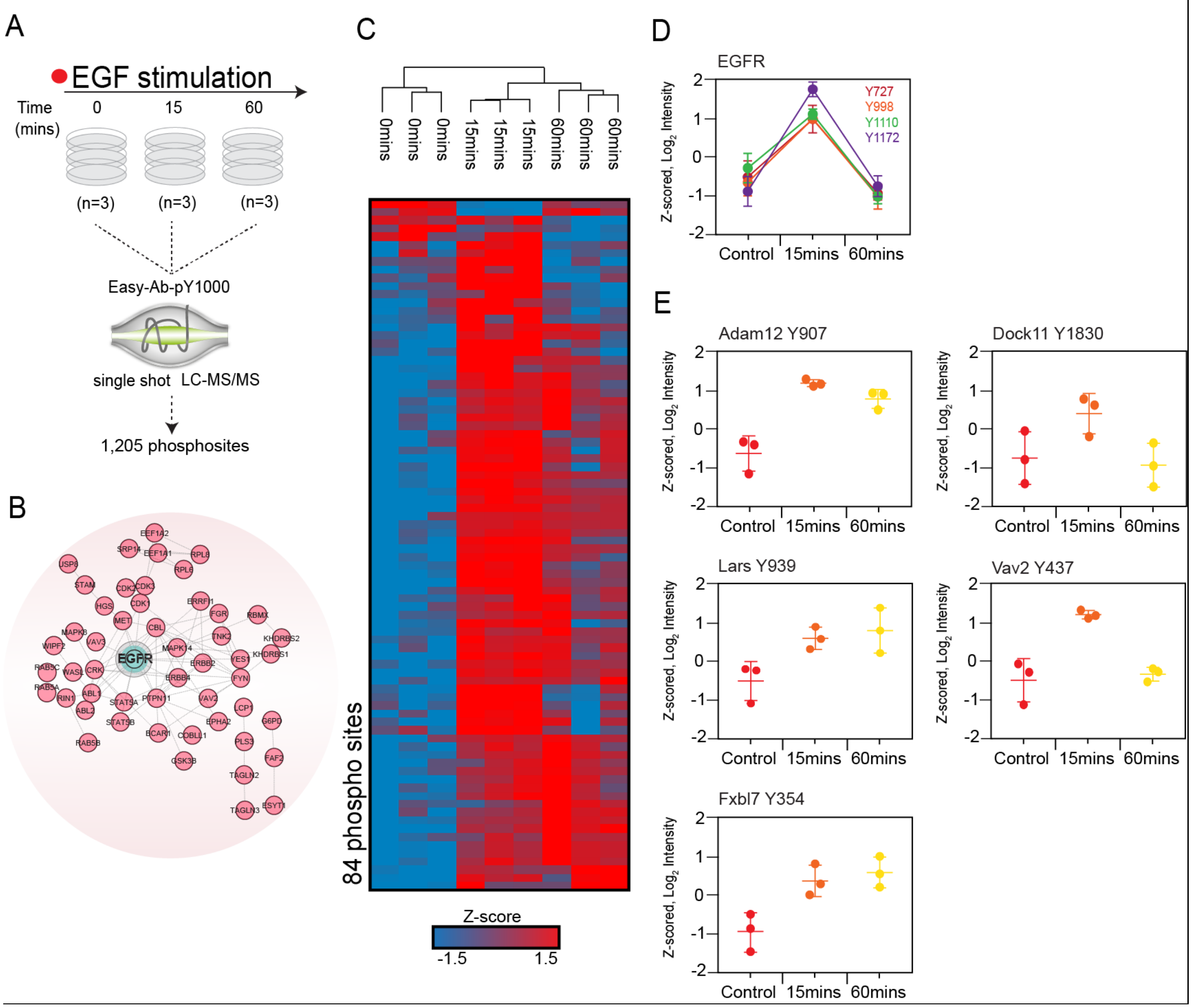


**Figure S4 EasyAb facilitates time resolved EGFR signaling.**

(**A**) Experimental set up of time resolved EGFR signaling. A single 15cm petri dish with 70% confluent HeLa cells per sample per condition were used. HeLa cells were stimulated by EGF at time points 0, 15 and 60 minutes in biological triplicates. Samples were processed as described in the materials and methods section and subjected to MS analysis.

(**B**) String network map of EGFR signaling. The significantly phosphorylated proteins upon EGF activation The EGFR node in turquoise colour and interactors node in pale red.

(**C**) Heatmap represents 84 regulated phosphosites that were temporally regulated upon EGF stimulation as illustrated.

(**D**) Profile plot shows the regulation of EGFR of phosphotyrosine sites presented as z-scored $\log_2$ phosphosite intensities.

(**E**) Profile plot shows novel EGFR signaling members activated upon EGF stimulation. The phosphotyrosine phosphorylation profile of each novel target varies with time. Data presented as z-scored $\log_2$ phosphosite intensities.

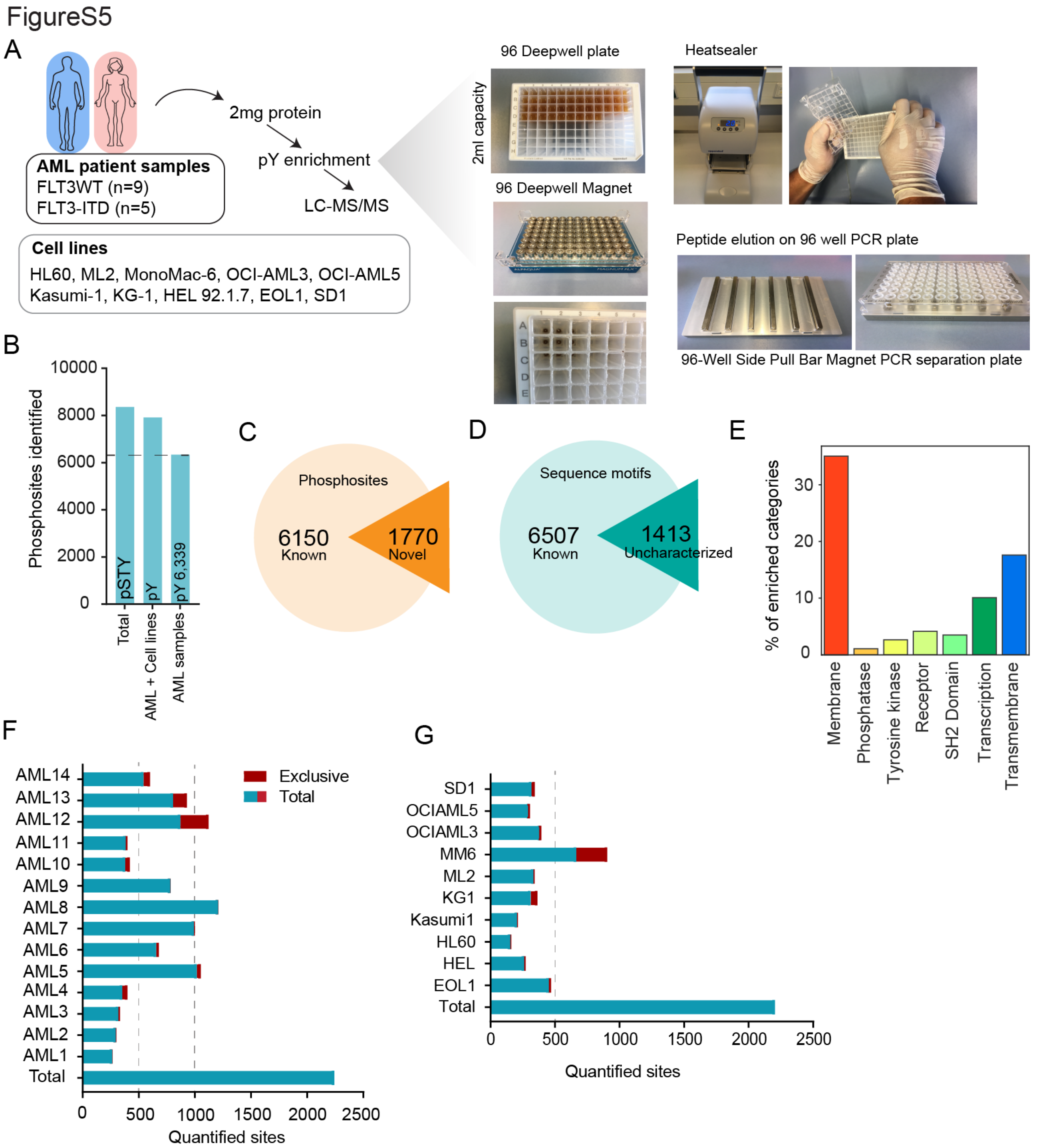


**Figure S5 EasyAb high-throughput workflow.**

(**A**) High throughput EasyAb setup optimized for parallel 96 sample processing. Details of the equipment are described in the method section. AML patient sample and in leukemic cell lines were used to test the application.
(**B**) Bar plot showing the identified phosphotyrosine sites in AML patient samples and cell lines.
(**C**) Pie chart shows the number of known and novel phosphotyrosine sites identified in the dataset.

(**D**) Pie chart shows the number of phosphotyrosine sites carrying characterized and uncharacterized sequence motif.
(**E**) Bar plot shows enrichment percentage of phosphotyrosine phosphorylated membrane and transmembrane proteins in the dataset.
(**F-G**) Total and exclusive number of phosphotyrosine sites quantified in (**E**) patient samples (**F**) cell lines.

FigureS6

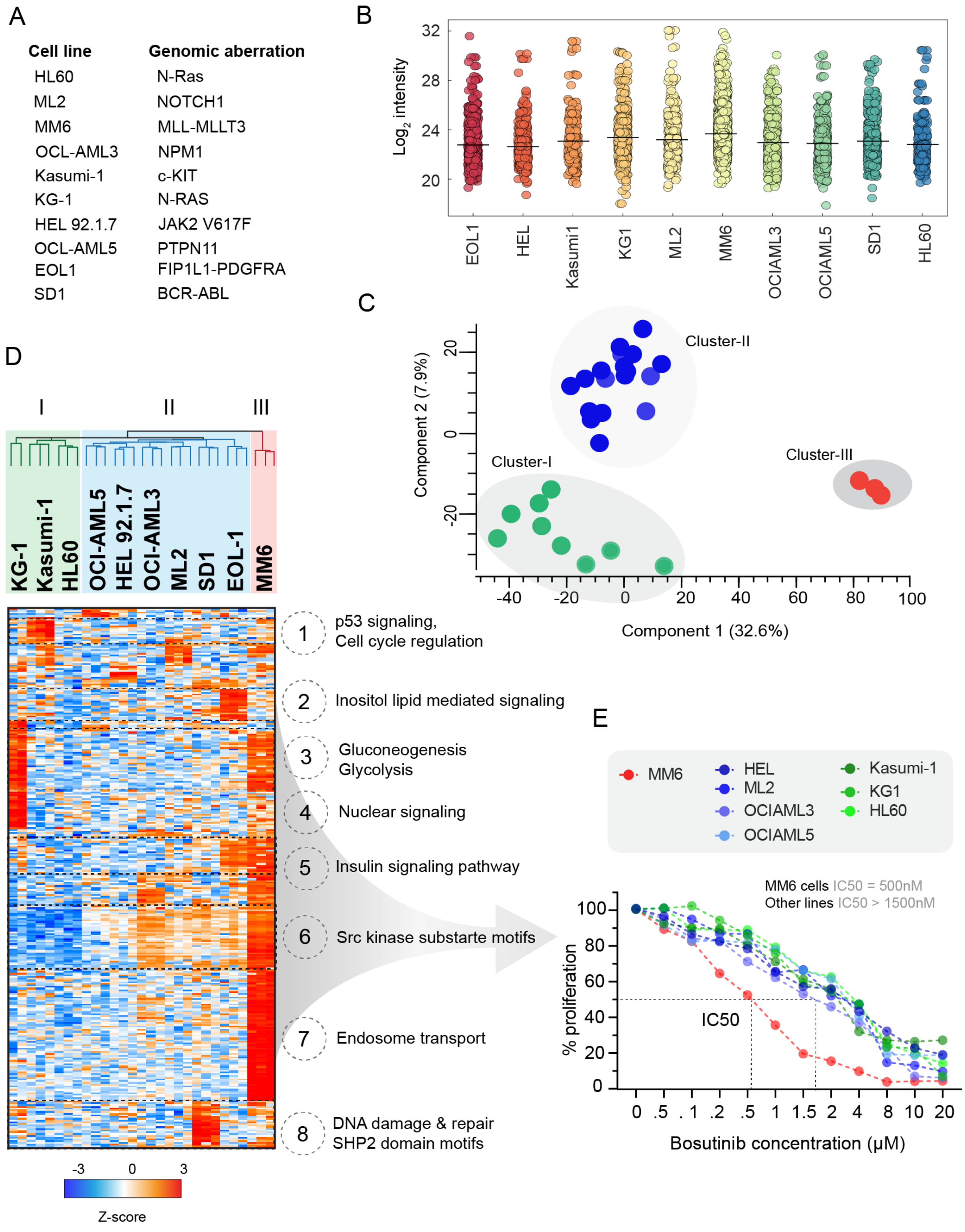
A
Cell line
Genomic aberration
HL60 N-Ras
ML2 NOTCH1
MM6 MLL-MLLT3
OCL-AML3 NPM1
Kasumi-1 c-KIT
KG-1 N-RAS
HEL 92.1.7 JAK2 V617F
OCL-AML5 PTPN11
EOL1 FIP1L1-PDGFRA
SD1 BCR-ABL
B
Log2 intensity
32
28
24
20
EOL1
HEL
Kasumi1
KG1
ML2
MM6
OCIAML3
OCIAML5
SD1
HL60
C
Cluster-II
Cluster-I
Cluster-III
Component 2 (7.9%)
Component 1 (32.6%)
D
I
II
III
KG-1
Kasumi-1
HL60
OCI-AML5
HEL 92.1.7
OCI-AML3
ML2
SD1
EOL-1
MM6
1 p53 signaling, Cell cycle regulation
2 Inositol lipid mediated signaling
3 Gluconeogenesis Glycolysis
4 Nuclear signaling
5 Insulin signaling pathway
6 Src kinase substarte motifs
7 Endosome transport
8 DNA damage & repair SHP2 domain motifs
-3 0 3
Z-score
E
MM6
HEL
ML2
OCIAML3
OCIAML5
Kasumi-1
KG1
HL60
MM6 cells IC50 = 500nM
Other lines IC50 > 1500nM
IC50
% proliferation
Bosutinib concentration (μM)

**Figure S6 Phosphotyrosine profiling of leukemia cell lines**

(**A**) List of AML cell lines with known driver gene mutations. The samples were processed in their native condition as biological triplicates in a parallel 96 deep well format for the enrichment of phosphophotyrosine peptides using 100µl of pY1000-magnetic bead conjugated antibody in 2mg of lysate. (**B**) The $log_2$ intensity of the phosphotyrosine sites quantified in each cell lines.

(**C**) Principle component analysis (PCA) clusters 10 cell lines into 3 groups based on the phosphotyrosine phosphoproteome.

(**D**) Unsupervised hierarchical cluster of ANOVA t-test significant phosphosites. Heatmap showing the enrichment of Src kinase substrate motifs and receptor signaling, confirming the presence of three major cell line groups matching the PCA data.

(**E**) Cell growth curve shows proliferation of cells in response to increasing concentration of Src kinase inhibitor Bosutinib measured by MTS proliferation assay. Cluster-III cell line MM6 displays highest sensitivity (IC50 = 500 nM) to Bosutinib compared to cluster-I and cluster-II cell lines. Data obtained from 2-3 independent experiment each with 8 technical replicates.

FigureS7

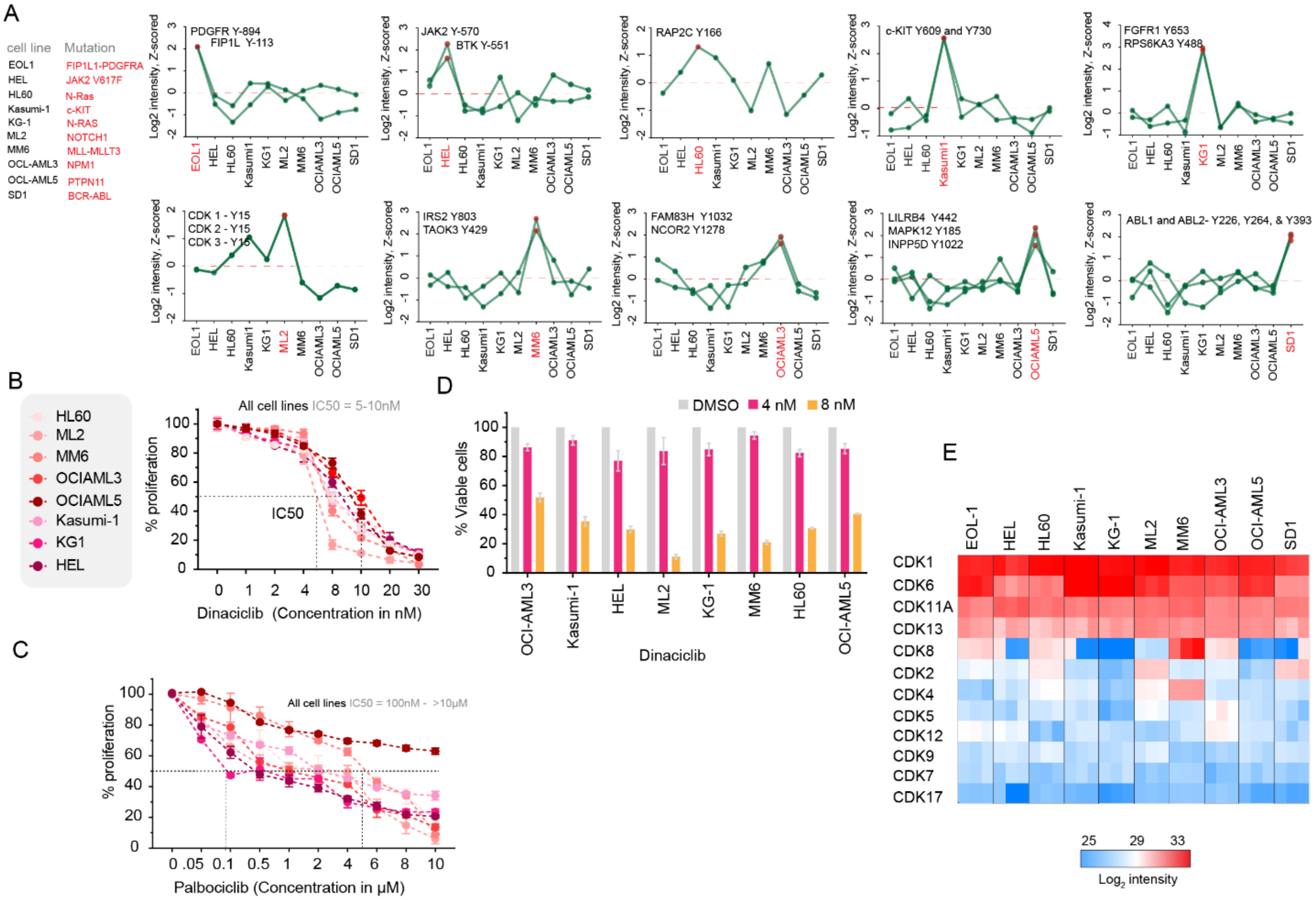


**Figure S7 AML cell lines specific phosphotyrosine signature and sensitivity to CDK 1/2/5/9 inhibitor Dinaciclib.**

(**A**) Cell line specific known and novel tyrosine phosphorylated protein profile presented as z-scored $log_2$ phosphosite intensities. Quantification of known cell line specific phosphotyrosine changes: PDGFR-FIP1L expressing EOL-1 cells with PDGFR Y894 and FIPL1 Y113, JAK2 Y570 in HEL cells, cKIT Y609 phosphorylation in Kasumi-1 cells, FGFR1 Y653 in KG1 cells and ABL Y226 in SD1 cells. Profile of cell line specific novel changes: RAP2C Y166 in HL60 cells, FAM83H Y1032 in OCIAML3, LILRB4 Y442 in OCIAML5, TAOK3 Y429 in MM6 cells, CDK1/2/3 Y15 in ML2 cells.
(**B**) Growth curve shows proliferation of cell lines in response to increasing concentration of CDK1/2/5/9 inhibitor Dinaciclib measured by MTS proliferation assay. ML2 display highest sensitivity (IC50 = ~ 5 nM) to Dinaciclib compared to OCIAML3 cell line which has the highest IC50 = ~10nM. Data obtained from 2-3 independent experiment each with 8 technical replicates and mean ± standard deviation.
(**C**) Growth curve shows proliferation of cell lines in response to increasing concentration of CDK4/6 inhibitor Palbociclib measured by MTS proliferation assay. Cell lines show IC50 between 100 nM- >10 μM. Data obtained from 2-3 independent experiment each with 8 technical replicates and mean ± standard deviation.
(**D**) Barplot displays the percentage of viable cells after 48 hours of Dinaciclib treatment at 4 nM and 8 nM measured by Trypan Blue. ML2 cells show >90% cell death compared to other cell lines at the dose of 8 nM.
(**E**) Heat map displays the expression of CDK proteins the AML cell lines. Data presented as $log_2$ protein intensity in biological triplicates.

FigureS8

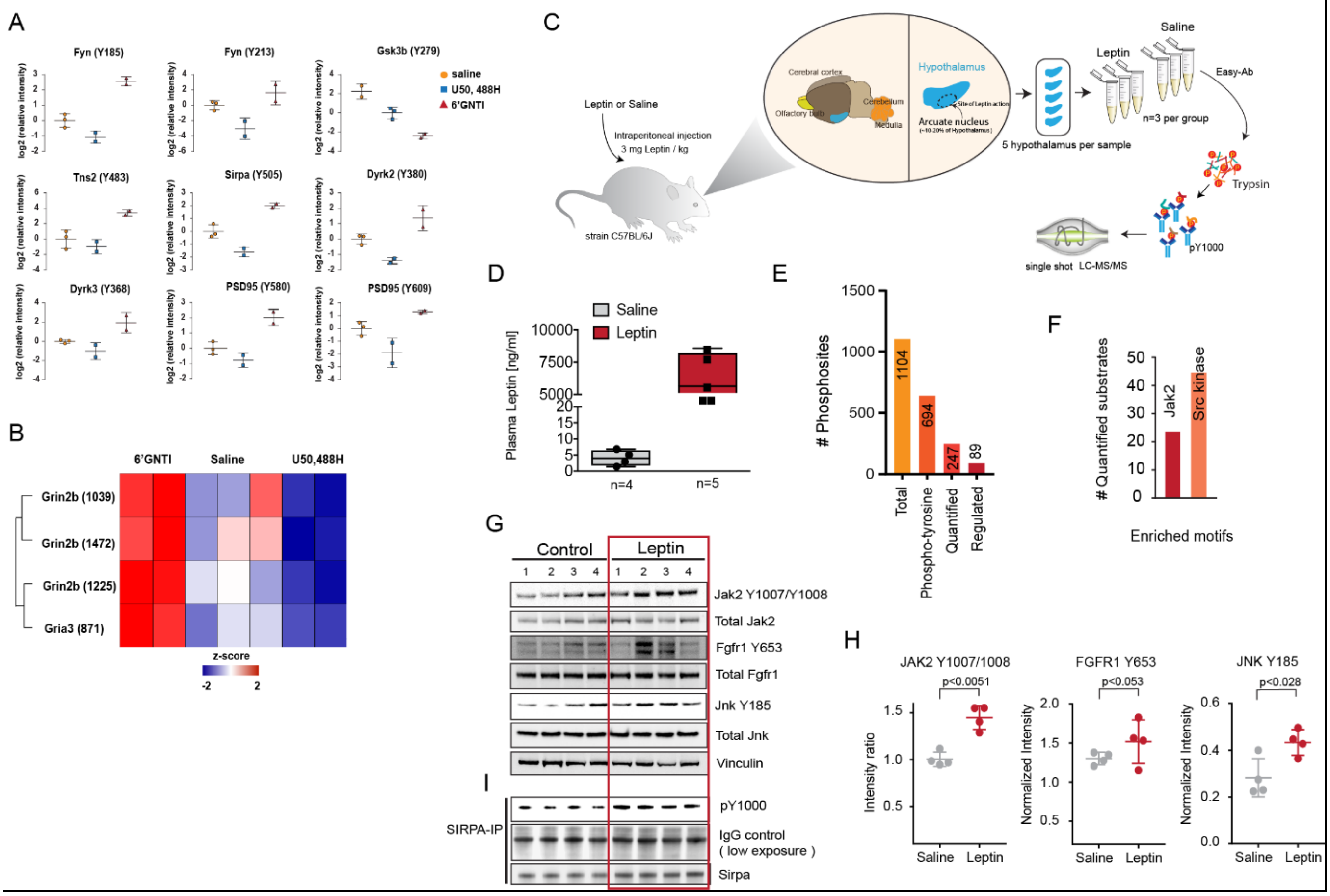


**Figure S8 In vivo GPCR and leptin induced phosphotyrosine signaling.**

(**A**) Selected ANOVA significant sites in the KOR study. Quantification of each measurement is transformed with log2 and normalized against the mean of their respective saline control. Mean and standard deviation of each condition is shown as interleaved bars.

(B) Phosphotyrosine sites on NMDA receptor subunit epsilon-2 (Grin2b) and subunit GluA3 of α-amino-3-hydroxy-5-methyl-4-isoxazolepropionic acid (AMPA) receptor (Gria3). The z-score of individual phosphosite is used for construction of the heatmap.

(**C**) Overview of leptin experiment. Saline or Leptin were injected intraperitoneally and mice sacrificed 30 minutes post treatment. Hypothalami were dissected, snap frozen, processed according to EasyAb protocol, and samples analyzed with single run LC-MS/MS.

(**D**) Box plot shows leptin levels in the plasma of leptin injected vs saline injected mice measured by ELISA test. On average leptin levels were 1000-fold higher in the leptin injected mice.

(**E**) Summary of identified, quantified and regulated phosphosites (student t-test p-value) in the dataset.

(**F**) Bar plot shows number of phosphosites enriched with Jak2 kinase and Src kinase substrate motifs. (**G**) Immuno blot displays the phosphorylation of JAK2 (antibody for autophosphorylation site Y570 were not available, therefore Y1007/1008 was utilized to assess the Jak2 activation in western blot), JNK, and FGFR1 upon saline (n = 4) and leptin (n = 4) treatment after 30 mins in mouse hypothalami.

(**H**) Scatter dot blot represents the intensity ratio of phosphorylated Jak2, Jnk and FGFR1 normalized to its total protein levels detected in saline and leptin samples. p-values calculated using pairwise two-tailed t-test are shown.

(**I**) Immunoprecipitation of SIRPA and detection of tyrosine phosphorylated SIRPA using phosphotyrosine antibody pY1000 in the hypothalami of saline (n = 4) and leptin (n = 4) treated mice.
